# Maladaptive myelination promotes epileptogenesis in absence epilepsy

**DOI:** 10.1101/2020.08.20.260083

**Authors:** Juliet K. Knowles, Caroline Soane, Eleanor Frost, Lydia T. Tam, Danielle Fraga, Haojun Xu, Ankita Batra, Lijun Ni, Katlin Villar, Tristan Saucedo, John Huguenard, Michelle Monje

## Abstract

Neuronal activity can influence the generation of new oligodendrocytes (oligodendrogenesis) and myelination. In health, this is an adaptive process that can increase synchrony within distributed neuronal networks and contribute to cognitive function. We hypothesized that in seizure disorders, aberrant neuronal activity may promote maladaptive myelination that contributes to pathogenesis. Absence epilepsy is a disease defined by increasingly frequent behavioral arrest seizures over time, thought to be due to thalamocortical network hypersynchrony. We tested the hypothesis that activity-dependent myelination resulting from absence seizures promotes epileptogenesis. Using two distinct models of absence epilepsy, Wag/Rij rats and *Scn8a*^+/mut^ mice, we found increased oligodendrogenesis and myelination specifically within the absence seizure network. These changes are evident only after seizure onset in both models and are prevented with pharmacological inhibition of seizures. Genetic blockade of activity-dependent myelination during epileptogenesis markedly decreased seizure frequency in the *Scn8a*^+/mut^ mouse model of absence epilepsy. Taken together, these findings indicate that activity-dependent myelination driven by absence seizures contributes to seizure kindling during epileptogenesis.

## Introduction

Neuronal activity can modulate myelin development (Makinodan et al., 2012; Hines et al., 2015; Mensch et al., 2015) and promote new oligodendrocyte generation and myelination in cortical and callosal axons throughout life (Liu et al., 2012; Gibson et al., 2014; Hughes et al., 2018; Mitew et al., 2018; Swire et al., 2019; Steadman et al., 2020). Activity-regulated myelination is adaptive in the healthy brain, increasing neural network synchrony (Pajevic et al., 2014; Noori et al., 2020; Steadman et al., 2020) and contributing to cognitive functions including learning, attention and memory consolidation (McKenzie et al., 2014; Xiao et al., 2016; Geraghty et al., 2019; Steadman et al., 2020). The effects of myelin plasticity on network function in health raises the question of how activity-regulated myelination may modulate network function in disease states characterized by abnormal patterns of neuronal activity, such as epilepsy. Diffusion tensor imaging (DTI) has demonstrated abnormal white matter microstructure in various forms of epilepsy including absence epilepsy in humans and rodent models; however, definitive conclusions cannot be drawn about underlying white matter structure, nor is it known how altered white matter structure may contribute to epilepsy pathophysiology (Chahboune et al., 2009; Gross, 2011; Yang et al., 2012; van Luijtelaar et al., 2013; Weiskopf et al., 2015).

Absence seizures occur in multiple forms of human epilepsy, and are associated with behavioral arrest and generalized but frontally predominant 3-4 Hz spike-wave discharges (Dlugos et al., 2013; Guilhoto, 2017). Seizures arise from abnormal oscillations between the thalamus and cortex and propagate along myelinated tracts including the anterior portions of the corpus callosum (Musgrave and Gloor, 1980; Vergnes et al., 1989; Holmes et al., 2004). In humans and rodents, absence seizures are brief but very frequent, occurring hundreds of times per day (Coenen and Van Luijtelaar, 1987). Thus, absence epilepsy presents an ideal paradigm to examine the relationship between activity-regulated myelination and seizure pathophysiology.

Rodent models of absence epilepsy exhibit defined periods of epileptogenesis in which seizures begin and then increase in daily frequency over time (Coenen and Van Luijtelaar, 1987; Dezsi et al., 2013; Makinson et al., 2017). This pattern of developmental seizure onset with rapid, progressive worsening over time reflects the natural history of untreated absence epilepsies in children (Brigo et al., 2018). Blockade of seizures during this window in one model of absence epilepsy, Wag/Rij rats, prevents or delays epileptogenesis (Blumenfeld et al., 2008; van Luijtelaar et al., 2013; Leo et al., 2019), indicating that seizure onset induces pathological network changes that contribute to subsequent progression in seizure frequency (kindling). While mechanisms of absence seizure kindling are incompletely understood, a common feature is excessive synchrony (coordinated firing of groups of neurons) in the thalamocortical network (Huntsman et al., 1999; Bai et al., 2011; Makinson et al., 2017; Tangwiriyasakul et al., 2018). Given the effect of activity-regulated myelination on network synchrony (Noori et al., 2020; Steadman et al., 2020), we hypothesized that abnormally increased myelination induced by seizures might contribute to increasing seizure frequency during epileptogenesis.

## Results

### Oligodendrogenesis increases within the absence seizure network after seizure onset

To test the putative relationship between absence seizures and myelination, we used a well-established model of absence epilepsy, Wag/Rij rats (Coenen and Van Luijtelaar, 1987; Blumenfeld et al., 2008; Russo et al., 2016; Sorokin et al., 2017; Citraro et al., 2020). Wag/Rij is an inbred rat strain that develops spontaneous, stereotyped absence seizures characterized by brief behavioral arrest, similar to the episodes that occur in children with absence epilepsy (Wirrell, 2003; Russo et al., 2016). The EEG correlate of these episodes in Wag/Rij rats are ∼8 Hz, generalized, frontally predominant spike-wave discharges that are maximal in the somatosensory cortices (Coenen and Van Luijtelaar, 2003; van Luijtelaar and Sitnikova, 2006). Absence seizures arise from connections between the thalamus and the cortex (Williams, 1953; Masterton et al., 2013; Tenney et al., 2013; McCafferty et al., 2018). In rodents, absence seizures are particularly prominent in relays between the ventrobasal nuclear complex of the thalamus and somatosensory cortex, driven by complex circuitry involving interneurons of the reticular thalamic nucleus (Kao and Coulter, 1997; Meeren et al., 2002; Fogerson and Huguenard, 2016; Makinson et al., 2017). Seizures propagate throughout the brain via myelinated tracts including the internal capsule (interconnects the thalamus and cortex) and the corpus callosum, a commissural tract which is required for seizure generalization (Musgrave and Gloor, 1980; Vergnes et al., 1989) (**Figure 1A**). Seizures in Wag/Rij rats develop over a well-defined period of epileptogenesis: infrequent seizures begin around 2 months of age and steadily increase in daily frequency until around 6 months of age, when the rate plateaus at 20-30 seizures per hour (Blumenfeld et al., 2008; van Luijtelaar et al., 2013). A closely related rat strain from which Wag/Rij is derived, Wistar, does not typically develop absence seizures during this time frame and therefore is used as a control for Wag/Rij (Blumenfeld et al., 2008; Chahboune et al., 2009; Sarkisova et al., 2010).

**Figure 1:**
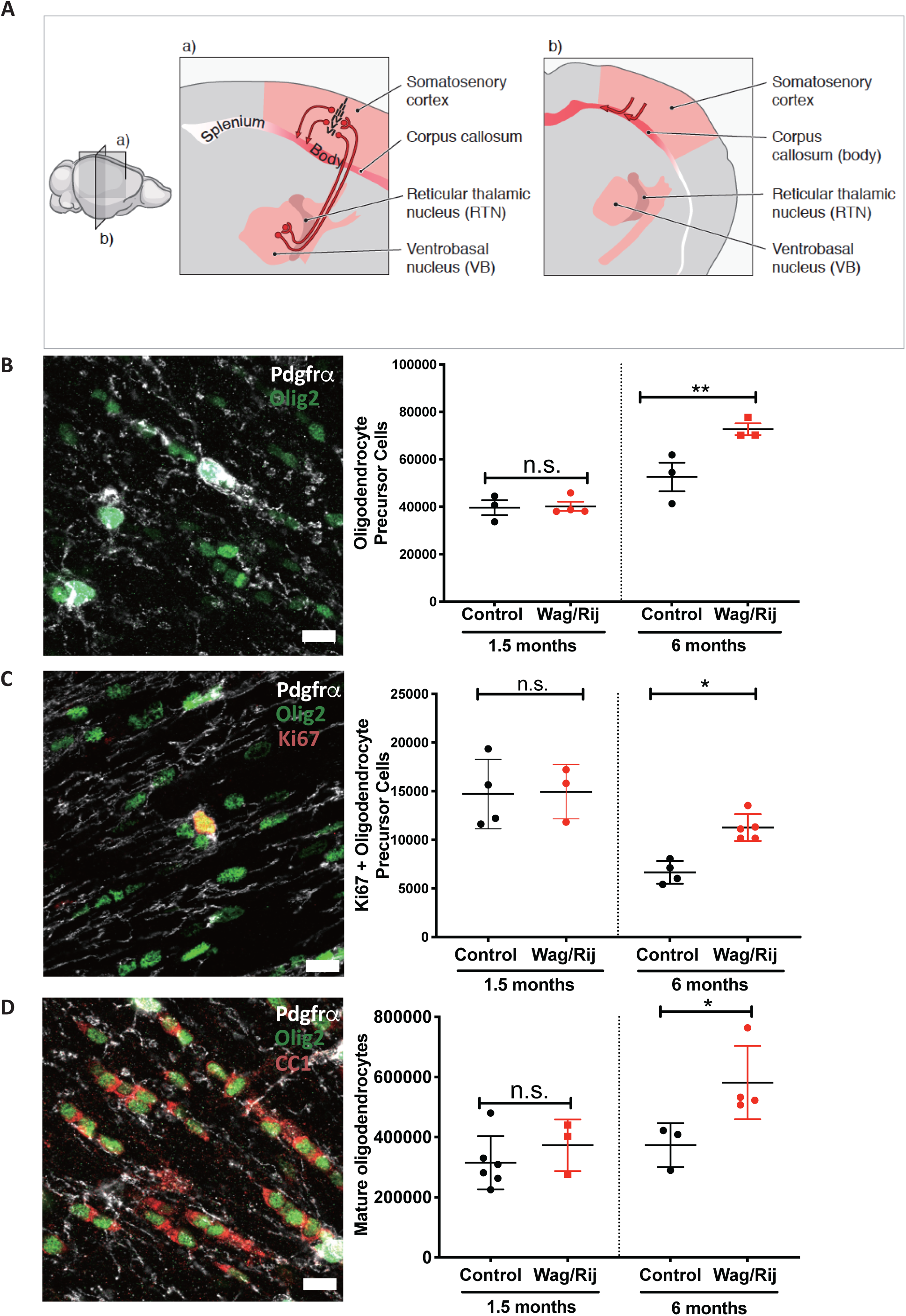
Oligodendrogenesis increases within the absence seizure network after seizure onset in Wag/Rij rats. **(A) Schematic of absence seizure network illustrating propagation through the body of the corpus callosum**. Schematic demonstrates sagittal (a) and coronal (b) views of the absence seizure network, shown in pink/red. Absence seizures result from hypersynchronous oscillations between the thalamus and cortex; in rodents, seizure activity is particularly prominent in connections between the ventrobasal and reticular thalamic nuclei and somatosensory cortices. Seizure activity propagates across the body of the corpus callosum, leading to bi-hemispheric generalization. In humans, absence seizures are frontally predominant and in rodents, there is little involvement of the occipital cortices and posterior region of the corpus callosum (splenium) that connects them. **(B) Absence epileptogenesis is associated with increased callosal oligodendrocyte precursor cells**. Left: representative photomicrograph of callosal oligodendrocyte progenitor cells (OPCs) from a 6-month old control rat co-expressing Olig2 (green) and PDGFR*α* (white); scale bar, 10 μm. Other oligodendroglial lineage cells express Olig2 (green only) but not PDGFR*α*. Right: Unbiased stereological quantification of oligodendrocyte precursor cells (OPCs) in the body of the corpus callosum at 1.5 months of age (prior to seizure onset) and 6 months of age (after seizures are well-established in Wag/Rij rats) in control (Wistar) and Wag/Rij rats. Black dots represent control rats and red dots represent Wag/Rij rats. Each data point represents total OPC number from 1 rat; 477-909 cells were counted per rat (1.5-month timepoint) and 735-1154 cells were counted per rat at the 6-month timepoint. Data represent mean ± SEM. 1.5-month timepoint, n = 3 control, 4 Wag/Rij rats; 6-month timepoint, n = 3 control, 3 Wag/Rij rats. **(C) Absence epileptogenesis is associated with increased callosal OPC proliferation**. Left: representative photomicrograph of a dividing OPC from a 6-month-old control rat co-expressing Olig2 (green), PDGFR*α* (white) and Ki67 (red). Scale bar is 10 μm. Right: Unbiased stereological quantification of proliferating oligodendrocyte precursor cells (OPCs) in the body of the corpus callosum at 1.5 months of age (prior to seizure onset) and 6 months of age (after seizures are well-established in Wag/Rij rats) in control (Wistar) and Wag/Rij rats. Each data point represents total Ki67-OPC number for one rat. At the 1.5-month timepoint, 426-734 cells were counted per rat, while 229-448 cells were counted per rat at the 6-month timepoint (1.5-month timepoint, n = 4 control, 3 Wag/Rij; 6-month timepoint, n = 4 control, 5 Wag/Rij rats). **(D) Absence epileptogenesis is associated with increased callosal oligodendrocytes**. Left: representative photomicrograph of mature oligodendrocytes in the corpus callosum of a 6-month old control rat, co-expressing Olig2 (green) and CC1 (red). These cells are distinct from precursor cells, which express PDGFR*α* (white) and Olig2. Scale bar is 10 μm. Right: Unbiased stereological quantification of mature oligodendrocytes in the body of the corpus callosum at 1.5 months of age (prior to seizure onset) and 6 months of age (after seizures are well-established in Wag/Rij rats) in control (Wistar) and Wag/Rij rats. Each data point represents total mature oligodendrocytes for 1 rat; at the 1.5-month timepoint, 478-1102 cells were counted for each rat while at the 6-month timepoint, 757-1522 cells were counted for each rat. (1.5-month timepoint, n = 6 control, 3 Wag/Rij; 6-month timepoint, n = 3 control, 4 Wag/Rij rats). For all panels in this figure, data were analyzed by ANOVA with post-hoc Sidak’s test (comparing groups within 1.5 month or 6-month timepoints), correcting for multiple comparisons. For all panels, *p<0.05, **p<0.01, ***p<0.001, n.s. = p>0.05.

To investigate whether absence seizures cause aberrant activity-regulated myelination within the seizure network, we began by assessing oligodendrocyte precursor cell (OPC) proliferation together with total numbers of OPCs and mature oligodendrocytes in the mid-region (body) of the corpus callosum, focusing specifically on the area interconnecting the somatosensory cortices that is involved in the absence seizure network. Given the anatomical differences between Wag/Rij and Wistar (control) rats (**Supplemental Figure 1A-B**), we utilized unbiased stereological methods to assess total cell numbers as well as volume of the corpus callosum and cell density. Prior to seizure onset, at 1.5 months of age, control rats and Wag/Rij (seizure) rats have equivalent numbers of callosal OPCs. However, at 6 months of age, when seizures are well established, we found that Wag/Rij rats exhibit a significant increase in OPC (cells co-expressing PDGFR*α* and Olig2) number and density (**Figure 1B and Supplemental Figure 1C**) as well as dividing (Ki67 positive) OPCs (**Figure 1C**). We next determined whether increased numbers of precursor cells were associated with increased quantities of callosal oligodendrocytes (CC1, Olig2-expressing cells) in the same region of the corpus callosum. Wag/Rij rats also exhibit increased oligodendrocytes (total number and cell density) at 6 months of age, following the period of epileptogenesis, indicative of increased oligodendrogenesis (**Figure 1D and Supplemental Figure 1D**). In contrast, Wag/Rij and control rats exhibit similar numbers of oligodendrocytes at 1.5-months of age, prior to seizure onset (**Figure 1D)**. Taken together, these data indicate that oligodendrogenesis increases within the seizure circuit in parallel with epileptogenesis in the Wag/Rij rat model of absence epilepsy.

### Myelination increases within corpus callosum regions affected by absence seizures

Given that epileptogenesis is associated with increased callosal oligodendrogenesis in Wag/Rij rats, we next investigated whether myelin structure is also altered. We utilized transmission electron microscopy to visualize cross sections of myelinated axons in the mid-sagittal plane of the body of the corpus callosum (**Figure 2A**), where oligodendrogenesis was assessed. We measured myelin sheath thickness per axon diameter, *g*-ratio (Gibson et al., 2014; Geraghty et al., 2019; Steadman et al., 2020) in 1.5- and 6-month old Wag/Rij rats and Wistar controls. We found an increase in mean myelin sheath thickness (decreased *g*-ratio) in 6-month-old Wag/Rij rats compared to controls (**Figure 2B, D**). This difference in myelin was not observed prior to seizure onset at 1.5 months (**Figure 2B, C)** and is not attributable to strain differences in axon diameter (**Supplemental Figure 2A**).

**Figure 2:**
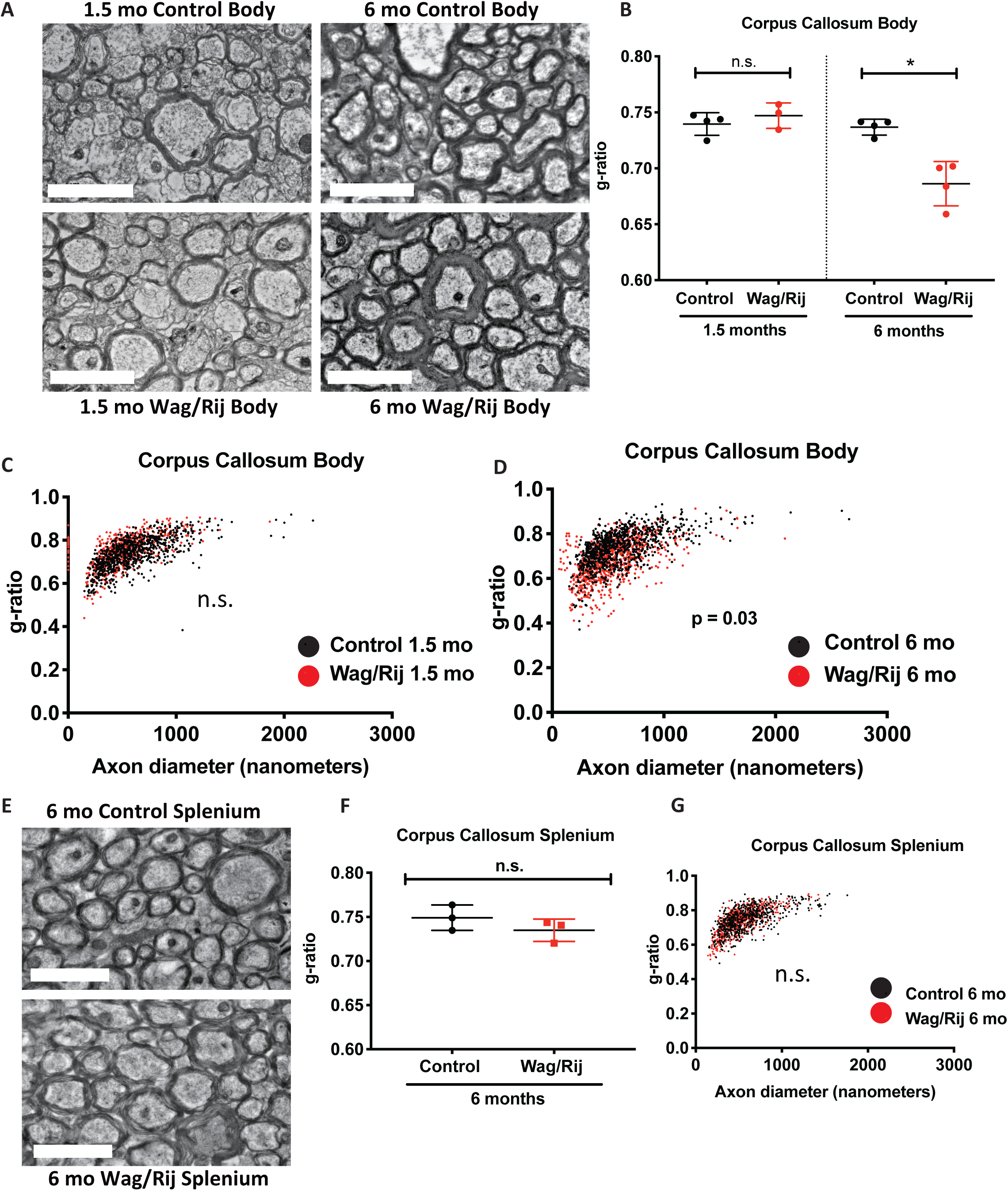
Myelination increases within the absence seizure network in Wag/Rij rats. **(A) Myelin sheaths appear thicker in Wag/Rij rats with seizures**. Representative transmission electron microscopy images of myelinated axon cross sections in the mid-sagittal body of the corpus callosum of 1.5-month (left) and 6-month-old (right) control rats (upper panels) and Wag/Rij rats (lower panels). Scale bar = 2 micrometers. **(B-D) Quantitative analysis indicates increased myelin sheath thickness in Wag/Rij rats after epileptogenesis. (B)** Mean *g*-ratio (axon diameter divided by the diameter of the entire fiber) of axons in the body of the corpus callosum in 1.5-month old and 6-month old Wag/Rij rats (red dots) and age-matched control rats (black dots). Smaller *g*-ratio values indicate thicker myelin sheaths. Each dot represents the mean *g*-ratio for one rat. For each rat, 195-264 axons were measured from 8-17 electron micrographs. Data represent mean ± SEM and were analyzed by ANOVA with post-hoc Sidak’s testing. 1.5-month timepoint, n = 4 control, 3 Wag/Rij rats; 6-month timepoint, n = 4 control, 4 Wag/Rij rats. **(C-D)** Scatterplots of individual axon *g*-ratio measurements which were shown as means in **(B)**, from 1.5-month-old rats **(C)** and 6-month-old rats **(D)**, as a function of axon diameter. Each data point represents one axon, with control axons in black and Wag/Rij axons in red. 1.5-month time-point: n = 4 control, 3 Wag/Rij rats; 6-month time-point: n = 4 control, 4 Wag/Rij rats. **(E-G) Increased myelination is specific to the seizure network. (E)** Representative transmission electron micrographs of myelinated axons in the splenium of a 6-month-old control rat (upper panel) and a Wag/Rij rat (lower panel). Scale bar = 2 μm. **(F)** Mean *g*-ratio of axons in the splenium of the corpus callosum in 6-month old Wag/Rij rats (red dots) and age-matched control rats (black dots). For each rat, 197-217 axons from 10-14 fields were quantified. n = 3 control, 3 Wag/Rij rats. Data represent mean ± SEM and were analyzed with a t-test. **(G)** Scatterplot of individual axon *g*-ratio measurements shown as means in **(F)**. Each data point represents one axon, with control axons in black and Wag/Rij axons in red. n = 3 control, 3 Wag/Rij rats. For all panels in this figure, *p<0.05, **p<0.01, ***p<0.001, n.s. = p>0.05.

Absence seizures in rodents are most prominent in the somatosensory cortices (Meeren et al., 2002; Polack et al., 2007; Scicchitano et al., 2015). We reasoned that if abnormally increased myelination is caused by seizure activity, these changes would be specific to the seizure-affected regions. We therefore assessed myelin in the posterior corpus callosum (splenium), connecting cortical regions where seizure activity is less prominent in humans and rodents (Meeren et al., 2002; Nersesyan et al., 2004; Moeller et al., 2010; Tenney et al., 2013; Meyer et al., 2018). The seizure-associated myelin difference observed in the body of the corpus callosum was not found in the splenium; **Figure 2E-G**. Taken together, these data indicate that seizures are associated with increased oligodendrogenesis and abnormally increased myelination in an anatomical pattern that parallels seizure activity.

### Seizures are necessary for aberrant callosal myelination

The temporal association between epileptogenesis and abnormally increased oligodendrogenesis and myelination suggests that seizures may induce aberrant activity-regulated myelination in Wag/Rij rats. In order to determine whether seizures are required for the observed increases in oligodendrogenesis and myelination, we treated Wag/Rij and control rats with the anti-seizure drug ethosuximide (ETX) at ∼300 mg/kg/day, a dose known to prevent or reduce seizures in Wag/Rij rats (Blumenfeld et al., 2008; Sarkisova et al., 2010). Similar to published work, this led to a mean plasma concentration of 101.3 ± 10.33 micrograms per mL (mean ± SEM, n= 20 rats), without signs of toxicity (Blumenfeld et al., 2008) and similar to therapeutic levels in humans, typically between 40-100 micrograms per mL (https://pubchem.ncbi.nlm.nih.gov/compound/Ethosuximide). Treatment was initiated at 1.5 months of age, prior to seizure onset. Following 5 months of treatment, EEG at 6.5 months of age revealed frequent absence seizures in vehicle-treated Wag/Rij rats (**Figure 3A-B**). Consistent with prior published data (Blumenfeld et al., 2008), mean seizure duration in vehicle-treated Wag/Rij rats was 5 ± 0.5 seconds (mean ± SEM, n=6 rats). As expected, treatment with ETX over the period of epileptogenesis significantly decreased or prevented seizures (**Figure 3B**).

**Figure 3:**
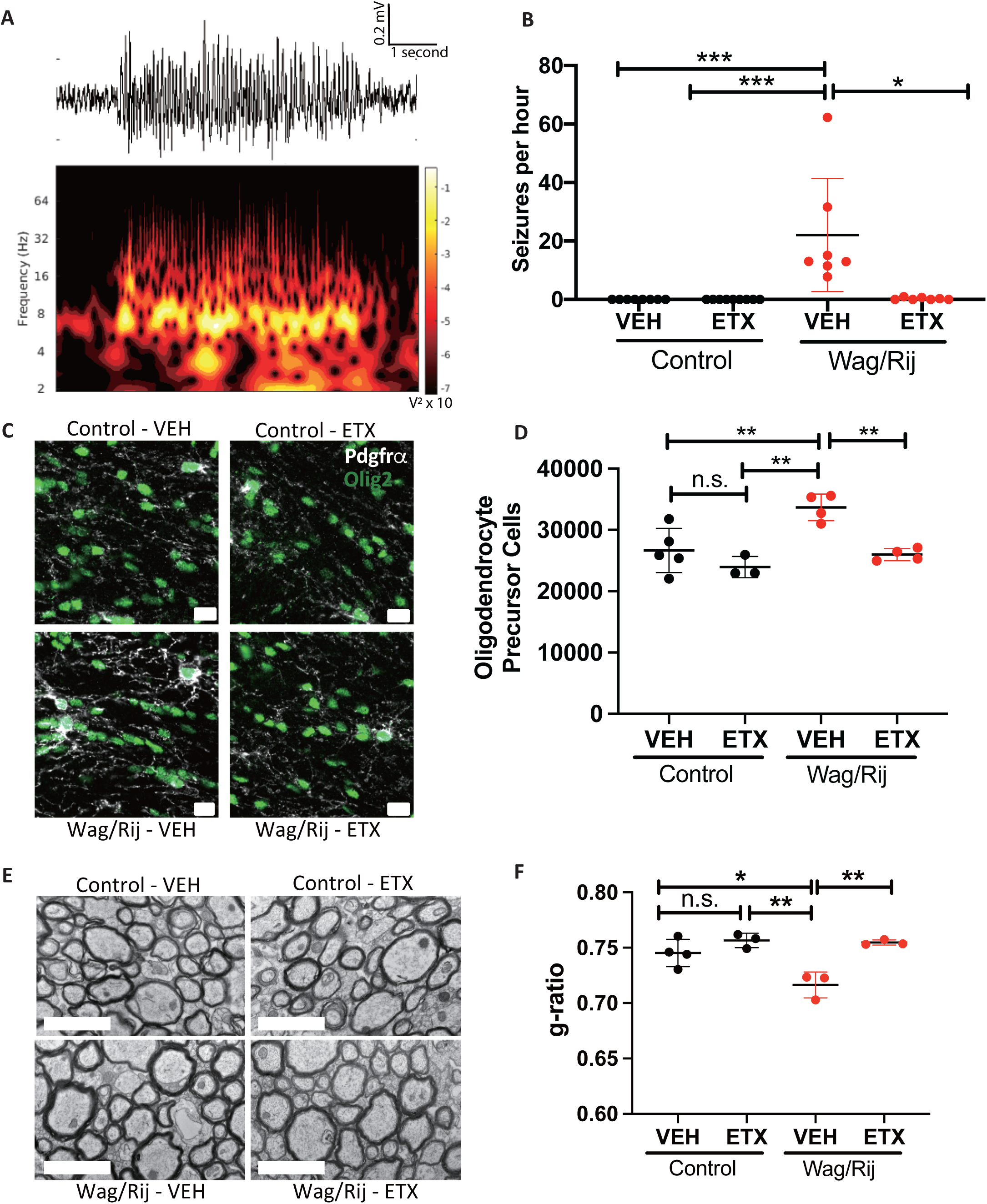
Seizures are necessary for aberrant callosal myelination. Control and Wag/Rij rats were treated with vehicle (VEH) or ethosuximide (ETX) during the period of epileptogenesis, from 1.5 months to 7 months of age. Electroencephalograms (EEGs) were recorded at 6.5 months of age. **(A)** Representative spike-wave discharge seizure from a 6.5-month-old vehicle-treated Wag/Rij rat (upper panel); spectral analysis demonstrating the predominant seizure frequency is 8 Hz (lower panel). **(B) Ethosuximide (ETX) decreases seizures in Wag/Rij rats**. Quantitative analysis of EEG recordings demonstrated well-established seizures in 6.5-month old vehicle (VEH)-treated Wag/Rij rats (22.0 ± 7.3 seizures per hour), whereas seizures were significantly decreased or absent in Wag/Rij rats treated with ETX (0.55 ± 0.31 seizures/hour). Each data point represents the mean seizures per hour for one rat; Control-VEH, n = 8 rats; Control-ETX, n = 9 rats; Wag/Rij-VEH, n = 7 rats; Wag/Rij-ETX, n = 7 rats. Data represent mean ± SEM and were analyzed by the Kruskal-Wallis and Dunn’s multiple comparisons tests. **(C-D) ETX treatment normalizes OPC number in Wag/Rij rats. (C)** Representative photomicrographs demonstrating increased OPCs (co-expressing PDGFR*α*, white and Olig2, green) in the body of the corpus callosum of 7-month old VEH-treated Wag/Rij rats compared to age-matched control rats treated with vehicle (VEH) or ETX and Wag/Rij rats treated with ETX. Scale bar = 10 μm. **(D)** Unbiased stereological quantification of OPCs in the body of the corpus callosum at 7 months of age in VEH or ETX-treated Wag/Rij or control rats. Each data point represents the OPC number for one rat; 474-926 cells were counted per rat. Note one half of the brain was used for these analyses and accordingly total OPC number measurements were ∼50% of those in Figure 1B, which utilized both sides of the brain. Control-VEH, n = 5 rats; Control-ETX, n = 3 rats; Wag/Rij-VEH, n = 4 rats; Wag/Rij-ETX, n = 4 rats. Data represent mean ± SEM and were analyzed by ANOVA with Tukey-Kramer post-hoc testing. **(E-F) ETX treatment normalizes myelination in Wag/Rij rats. (E)** Representative transmission electron micrographs demonstrating increased myelin sheath thickness in some axons in the body of the corpus callosum of 7-month old VEH-treated Wag/Rij rats compared to age-matched control rats treated with VEH or ETX and Wag/Rij rats treated with ETX. Scale bar = 2 μm. **(F)** Mean *g*-ratio of axons in the body of the corpus callosum in 7-month-old Wag/Rij (red dots) and control rats (black dots) treated with VEH or ETX, as determined by transmission electron microscopy. Each data point represents the mean *g*-ratio from 1 rat; 184-284 axons from 8-15 fields were quantified for each rat. Control-VEH, n = 4 rats; Control-ETX, n = 3 rats; Wag/Rij-VEH, n = 3 rats; Wag/Rij-ETX, n = 3 rats. Data represent mean ± SEM and were analyzed by ANOVA with Tukey-Kramer post-hoc testing. For all panels in this figure, *p<0.05, **p<0.01, ***p<0.001, n.s. = P>0.05.

We examined callosal OPC number and myelination from control or Wag/Rij hemi-brains following vehicle or ETX administration at 7 months of age. Similar to the findings described above (**Figures 1, 2**), OPC number and myelin sheath thickness were increased in vehicle-treated 7-month-old Wag/Rij rats compared to controls. However, ETX treatment normalized OPC number and myelin sheath thickness (*g-*ratio) in Wag/Rij rats (**Figure 3C-F**). ETX did not influence axonal diameter (**Supplemental Figure 2B**).

Together, these findings indicate that seizures increase myelination specifically within the seizure-affected region and suggest a mechanism of aberrantly increased activity-dependent myelination that could be deleterious (maladaptive), contributing to epilepsy pathogenesis. To further test this hypothesis, we sought to evaluate seizure-related myelin changes in a second model of absence epilepsy.

### Increased oligodendrogenesis and myelination in *Scn8a*^+/mut^ mice

We next quantified oligodendrogenesis and myelin structure in a second, distinct rodent model of absence epilepsy, *Scn8a*^+/mut^ mice. The use of this mouse model confers the advantage of wild-type littermates on a congenic background, and the opportunity for targeted genetic manipulation of activity-dependent myelination. *Scn8a*^+/mut^ mice bear a heterozygous loss of function mutation in the voltage-gated sodium channel *Nav1*.*6*, which results in abnormal thalamocortical hyper-synchrony and spontaneous 4-8 Hz absence seizures (Papale et al., 2009; Makinson et al., 2017). *Scn8a*^+/mut^ mice exhibit seizures that begin around post-natal day (P)21 and steadily increase in frequency until they occur ∼20-30 times per hour, by P35-P45 (Makinson et al., 2017).

We assessed callosal OPC proliferation and number in *Scn8a*^+/mut^ mice and littermate wild-type control mice (*Scn8a*^+/+^) before (P21) and after (P45) seizures are well established. Prior to seizure onset, callosal OPC proliferation and total number of callosal OPCs were equivalent. In contrast, after seizures are well established at P45, we found increased overall numbers of OPCs and proliferating OPCs in the corpus callosum of *Scn8a*^+/mut^ animals relative to littermate controls (**Figure 4A-D**). While corpus callosum volume was equivalent in *Scn8a*^+/mut^ and littermate control mice at P21, the volume of the corpus callosum was increased in *Scn8a*^+/mut^ mice at P45 (**Supplemental Figure 3A**), Concordant with previous work demonstrating constant density of OPCs throughout the murine brain (Hughes et al., 2013), we found that OPC density was similar in both groups when normalized to corpus callosum volume (**Supplemental Figure 3B**). The total number and callosal density of mature oligodendrocytes were found to be equivalent at P21, but increased in *Scn8a*^+/mut^ mice relative to *Scn8a*^+/^ at P45, indicative of increased oligodendrogenesis after seizure onset (**Figure 4E-F, Supplemental Figure 3C**).

**Figure 4:**
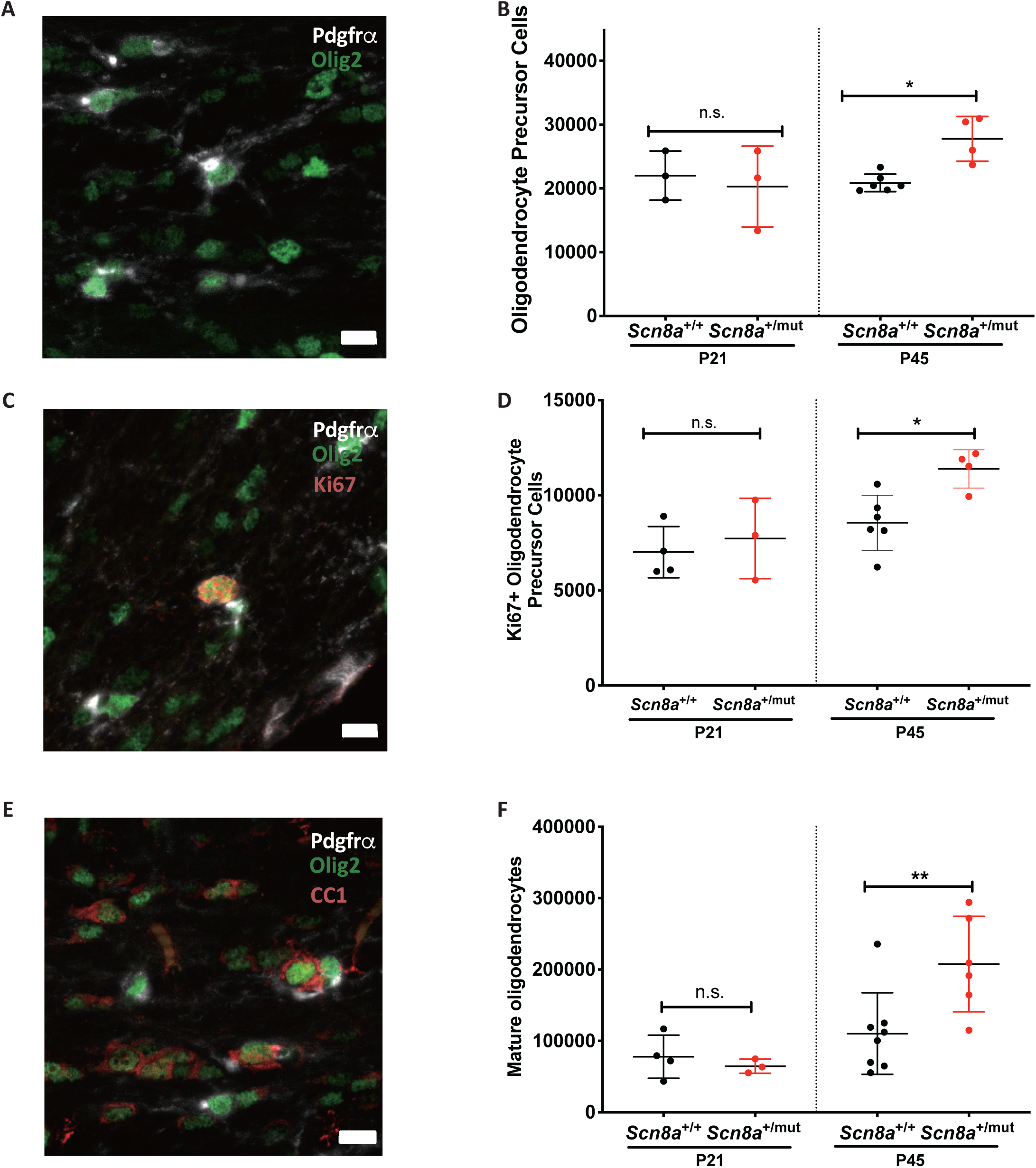
Increased oligodendrogenesis within the seizure network after seizure onset in. ***Scn8a***^***+/mut***^ **mice**. **(A-B) Absence epileptogenesis in *Scn8a***^***+/mut***^ **mice is associated with increased callosal OPC number. (A)** Representative photomicrograph of OPCs co-expressing PDGFR*α* (white) and Olig2 (green) from a *Scn8a*^+/+^ mouse. Other oligodendroglial lineage cells which are not OPCs express Olig2 but not PDGFR*α*. Scale bar = 10 μm. **(B)** Unbiased stereological quantification of OPCs in the body of the corpus callosum at 21 post-natal days (P21) (prior to seizure onset) and P45 (after seizures are well-established in *Scn8a*^+/mut^ mice). (P21, *Scn8a*^+/+^ n= 3 mice; *Scn8a*^+/mut^ n = 3 mice. P45, *Scn8a*^+/+^ n= 6 mice; *Scn8a*^+/mut^ n = 4 mice). Black dots represent wildtype littermates (*Scn8a*^+/+^) while red dots represent *Scn8a*^+/mut^ mice. Data represent mean ± SEM; each dot represents one animal. For each mouse, 348-672 cells (P21 mice) or 271-535 cells (P45 mice) were counted. **(C-D) Absence epileptogenesis in *Scn8a***^***+/mut***^ **mice is associated with increased OPC proliferation. (C)** Representative photomicrograph of dividing callosal OPC co-expressing Ki67 (red), PDGFR*α* (white) and Olig2 (green) from a *Scn8a*^+/+^ mouse. Scale bar = 10 μm. **(D)** Unbiased stereological quantification of proliferating OPCs in the body of the corpus callosum at P21 and P45 in *Scn8a*^+/mut^ (red dots) and *Scn8a*^+/+^ mice (black dots). Data represent mean ± SEM; each dot represents the number of proliferating OPCs for one animal. For each P21 mouse, 220-316 Ki67+ OPCs were counted; for each P45 mouse, 174-347 Ki67+ OPCs were counted. (P21, *Scn8a*^+/+^ n= 4 mice; *Scn8a*^+/mut^ n = 3 mice. P45, *Scn8a*^+/+^ n= 6 mice; *Scn8a*^+/mut^ n = 4 mice). **(E-F) Absence epileptogenesis in *Scn8a***^***+/mut***^ **mice is associated with increased oligodendrogenesis. (E)** Representative photomicrograph of callosal mature oligodendrocyte expressing CC1 (red) and Olig2 (green), but not PDGFR*α* (white) from a *Scn8a*^+/+^ mouse. Scale bar = 10 μm. **(F)** Unbiased stereological quantification of mature oligodendrocytes in the body of the corpus callosum at P21 (prior to seizure onset) and P45 (after seizures are well-established in *Scn8a*^+/mut^ mice) in *Scn8a*^+/mut^ and *Scn8a*^+/+^ mice. Data represent mean ± SEM; each dot represents one animal. For each P21 mouse, 380 – 748 mature oligodendrocytes were counted; for each P45 mouse, 555-2226 mature oligodendrocytes were counted (P21, *Scn8a*^+/+^ n= 4 mice; *Scn8a*^+/mut^ n = 3 mice. P45, *Scn8a*^+/+^ n= 8 mice; *Scn8a*^+/mut^ n = 6 mice). For all panels in this figure, data were analyzed with ANOVA with post-hoc Sidak’s test, correcting for multiple comparisons. *p<0.05, **p<0.01, ***p<0.001, n.s. = p>0.05.

The observed increase in corpus callosum volume after seizure onset (**Supplemental Figure 3A**) could be consistent with increased myelination. To determine whether increased oligodendrogenesis in *Scn8a*^+/mut^ mice is associated with increased myelination, we again assessed myelin structure in the midline sagittal body of the corpus callosum (**Figure 5**). This revealed that myelin sheath thickness was increased after seizure onset at P45 in *Scn8a*^+/mut^ mice relative to *Scn8a*^+/+^ littermate controls (**Figure 5A-B, D**). Prior to seizure onset at P21, *g*-ratios were equivalent in *Scn8a*^+/mut^ and littermate control mice. **(Figure 5A-C)**. Mean myelinated axon diameter was equivalent at P21 and at P45 in *Scn8a*^+/mut^ mice relative to *Scn8a*^+/+^ littermate controls, indicating that altered axon size does not contribute to *g*-ratio differences **(Figure 5F)**. Normalizing to differences in callosal volume, myelinated axon number was increased in P45 *Scn8a*^+/mut^ mice compared to *Scn8a*^+/+^ littermate controls. This difference in myelinated axons was not present prior to epileptogenesis at P21 **(Figure 5E)**.

**Figure 5:**
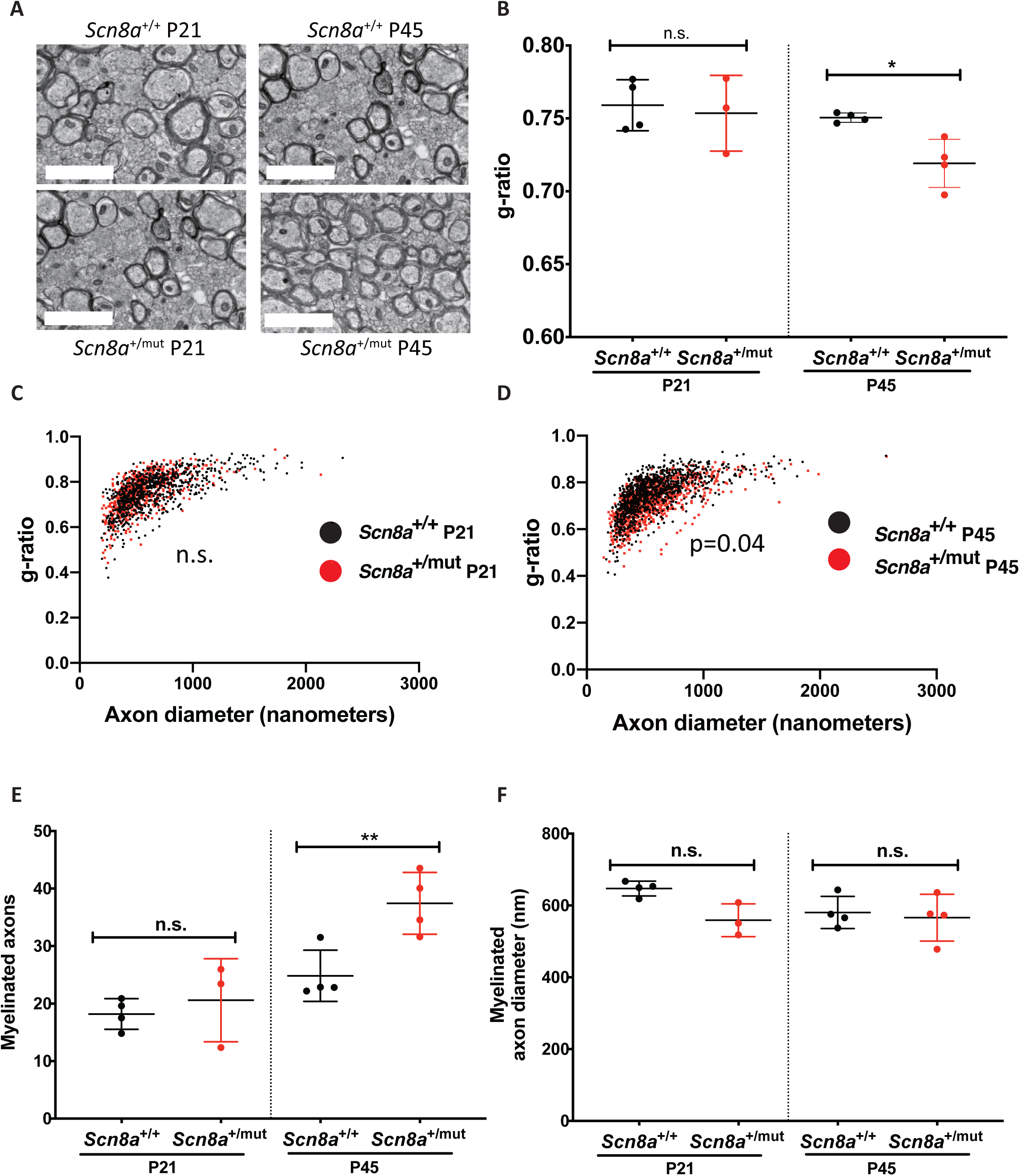
Increased callosal myelination after seizure onset in *Scn8a*^+/mut^ mice. **(A)** Representative transmission electron microscopy images of myelinated axons in the body of the corpus callosum of P21 (left) and P45 (right) *Scn8a*^+/+^ (upper panels) and *Scn8a*^+/mut^ mice (lower panels). Scale bar = 2 micrometers. **(B)** Quantitative analysis of myelin sheath thickness (*g*-ratio) in the body of the corpus callosum in at P21 (prior to seizure onset) and P45 (after seizures are well-established in *Scn8a*^+/mut^ mice), in *Scn8a*^+/mut^ (red dots) and *Scn8a*^+/+^ mice (black dots). Data represent mean ± SEM; each dot represents the mean *g*-ratio for one animal. For each mouse, 145-298 axons were quantified from at least 15 fields. (P21, *Scn8a*^+/+^ n= 4 mice; *Scn8a*^+/mut^ n = 3 mice. P45, *Scn8a*^+/+^ n= 4 mice; *Scn8a*^+/mut^ n = 4 mice). Data were analyzed by ANOVA with post-hoc Sidak’s test. **(C-D)** Scatterplot of individual axon *g*-ratio measurements from P21 **(C)** and P45 **(D)** mice, as a function of axon diameter, as in (**B)**. Each data point represents one axon, with *Scn8a*^+/+^ axons in black and *Scn8a*^+/mut^ axons in red. (P21, *Scn8a*^+/+^ n= 4 mice; *Scn8a*^+/mut^ n = 3 mice. P45, *Scn8a*^+/+^ n= 4 mice; *Scn8a*^+/mut^ n = 4 mice). **(E)** Myelinated axon number was quantified using transmission electron microscopy, in the body of the corpus callosum in P21 mice (prior to seizure onset) and P45 mice (with established seizures). Myelinated axon number was normalized to corpus callosum volume (**Supplemental Figure 3**). Each data point represents mean myelinated axons for one mouse, with black dots indicating *Scn8a*^+/+^ and red dots indicating *Scn8a*^+/mut^. For each mouse, axon number was quantified in 10 separate fields. Data represent mean ± SEM; data were analyzed with ANOVA followed by Sidak’s testing. (P21, *Scn8a*^+/+^=4 mice; *Scn8*^+/mut^ = 3 mice. P45, *Scn8a*^+/+^ = 4 mice; *Scn8a*^+/mut^ = 4 mice). **(F)** Myelinated axon diameters were quantified from transmission electron micrographs; black dots indicate *Scn8a*^+/+^ and red dots indicate *Scn8a*^+/mut^. Each dot represents the mean myelinated axon diameter from one mouse, with mean ± SEM indicated. Data were analyzed with ANOVA followed by Sidak’s testing. (P21, *Scn8a*^+/+^ n=4 mice; *Scn8a*^+/mut^ n = 3 mice; P45, *Scn8a*^+/+^ n= 4 mice, *Scn8a*^+/mut^ n = 4 mice.) For all panels in this figure, *p<0.05, **p<0.01, ***p<0.001, n.s. = p>0.05.

Taken together, findings in both Wag/Rij rat and *Scn8a*^+/mut^ mouse models demonstrate that absence seizures induce abnormally increased myelination within the affected thalamocortical seizure network. We next sought to determine the functional impact of seizure-associated myelination and tested the hypothesis that aberrantly increased myelination contributes to disease pathogenesis.

### Activity-dependent myelination contributes to epileptogenesis

In the healthy brain, activity-dependent myelination functions to synchronize regions within distributed neuronal networks, and this process is required for multiple forms of learning (Gibson et al., 2014; McKenzie et al., 2014; Xiao et al., 2016; Geraghty et al., 2019; Noori et al., 2020; Pan et al., 2020; Steadman et al., 2020). We hypothesized that seizure-associated, abnormally increased myelination might contribute to thalamocortical network hypersynchrony during epileptogenesis (Makinson et al., 2017), increasing disease severity. To assess the functional impact of myelin plasticity in absence epilepsy, we sought to block activity-dependent myelination during the period of epileptogenesis. We recently demonstrated that activity-dependent secretion of Brain Derived Neurotrophic Factor (BDNF) (Balkowiec and Katz, 2000; Hartmann et al., 2001; Dieni et al., 2012), and its subsequent signaling through the TrkB receptor on OPCs, is required for activity-dependent myelination of corticocallosal projection neurons (Geraghty et al., 2019). Conditional deletion of TrkB from OPCs prevents activity-dependent myelination in the corpus callosum but does not alter homeostatic oligodendrogenesis nor lead to myelin loss (Geraghty et al., 2019).

To enable blockade of activity-dependent myelination during epileptogenesis in *Scn8a*^+/mut^ mice, we generated *Scn8a*^+/mut^ and *Scn8a*^+/+^ littermates with floxed TrkB receptors (*Scn8a*^+/mut^; *TrkB*^fl/fl^ and *Scn8a*^+/+^; *TrkB*^fl/fl^). These mice were crossed with mice that express Cre under the PDGFR*α* promoter, in OPCs (Hughes et al., 2013)), following the administration of tamoxifen (*TrkB*^fl/fl^; *PDGFRα::Cre-ER*). Induction of Cre in this model leads to TrkB deletion in about 80% of OPCs (Geraghty et al., 2019); leak of Cre expression is not found in neurons (Mount et al., 2019). This cross yielded *Scn8a*^+/mut^; *TrkB*^fl/fl^ and *Scn8a*^+/+^; *TrkB*^fl/fl^ mice with or without inducible Cre expression in OPCs (referred to as *Scn8a*^+/+^ and *Scn8a*^+/+^ OPC cKO; *Scn8a*^+/mut^ and *Scn8a*^+/mut^ OPC cKO, respectively). All mice were treated with Tamoxifen (100mg/kg IP) between post-natal days 21-23 to ensure that any differences between genotype groups do not represent differences in Tamoxifen treatment. Following Tamoxifen treatment, mice were implanted for electrocorticography (EEG) to monitor seizure frequency in each group.

It should be noted that the original *Scn8a*^*+/mut*^ mouse line is on a C3HeB/FeJ background, while *Scn8a*^*+/mut*^;*TrkB*^*fl/fl*^ mice have a mixed C3HeB/FeJ and C57/BL6 background. Background strain can influence the age and kinetics of seizure onset (Ferraro et al., 1999; Papandrea et al., 2009). We therefore determined the timeline of epileptogenesis in *Scn8a*^*+/mut*^ mice with this mixed background. In *Scn8a*^*+/mut*^ mice (mixed background) with intact activity-dependent myelination, 4-8 Hz absence seizures begin around P45 (**Figure 6A, C-D**). Seizures then increase steadily and occur 20-30 times per hour, on average, by 4-6 months of age (**Figure 6A**).

**Figure 6:**
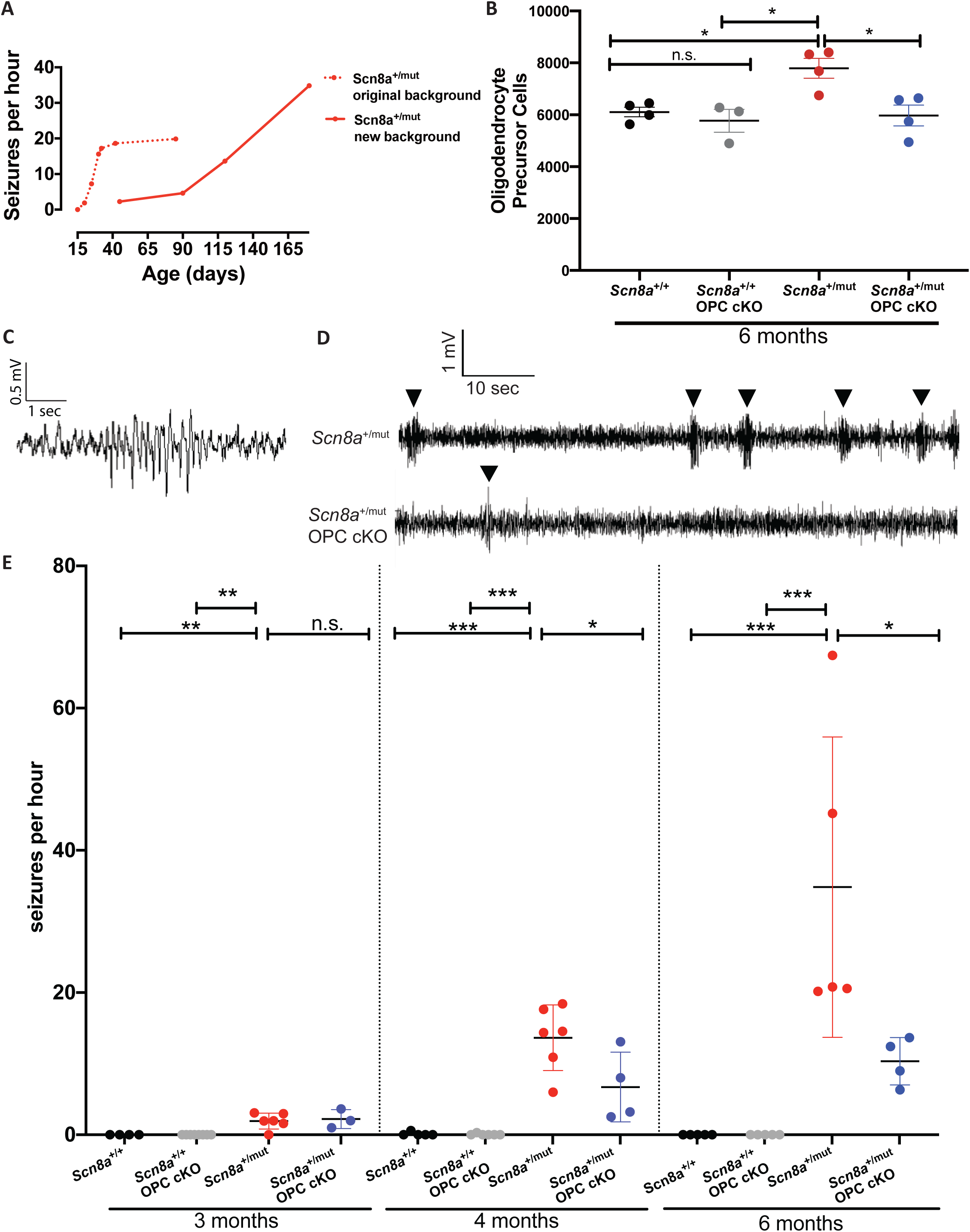
Activity-dependent myelination contributes to epileptogenesis. **(A) Epileptogenesis occurs later in *Scn8a***^**+/mut**^ **mice on a mixed genetic background**. In order to determine the role of activity-dependent myelination in epileptogenesis that occurs in *Scn8a*^+/mut^ mice (originally on a C3HeB/FeJ background), we generated *Scn8a*^+/mut^ mice with floxed TrkB receptor genes and the presence or absence of Cre under the PDGFR*α* promotor (*PDGFRα* - CreER). All mice underwent treatment with Tamoxifen; only mice expressing Cre subsequently underwent deletion of the TrkB receptor from OPCs. In mice with intact activity dependent myelination (*Scn8a*^+/mut^, *TrkB*^*fl/fl*^, *Cre* negative; C3HeB/FeJ and C57/BL6 mixed background; solid red line) we observed that seizure onset and progression occurred later time-points (P45-P180, i.e. 1.5 – 6 months), relative to the original *Scn8a*^+/mut^ line (on a C3HeB/FeJ background) in which seizures begin at ∼P21 and increase until P35-P45 (dashed red line, from Makinson et al, 2017). **(B) Genetic blockade of activity-dependent myelination prevents the oligodendroglial response to seizures**. Unbiased stereological assessment of OPC number (PDGFR*α*-expressing cells) in 6-month-old wild-type littermates with or without TrkB OPC expression (*Scn8a*^+/+^ and *Scn8a*^+/+^ OPC cKO, respectively), and *Scn8a*^+/mut^ and *Scn8a*^+/mut^ OPC cKO mice. Data represent mean ± SEM and were analyzed with ANOVA followed by Tukey-Kramer test, adjusting for multiple comparisons. Each dot represents one mouse; 123-273 OPCs were counted from hemi-brains for each mouse. *Scn8a*+/+ (black dots), n=4 mice; *Scn8a*^+/+^ OPC cKO (gray dots), n= 3 mice; *Scn8a*^+/mut^ (red dots), n=4 mice, *Scn8a*^+/mut^ OPC cKO (blue dots), n=4 mice. **(C-D)** Representative spike-wave discharge seizure in a *Scn8a*^+/mut^ mouse **©** and representative continuous EEG recordings from *Scn8a*^+/mut^ and *Scn8a*^+/mut^ OPC cKO mice **(D)**. **(E) Blockade of activity-dependent myelination decreases seizure frequency**. Quantitative analysis of seizure frequency from EEGs. Each data point represents mean seizures per hour for one mouse, show with group mean ± SEM. 3 month old time-point: *Scn8a*+/+ (black dots), n=4 mice; *Scn8a*^+/+^ OPC cKO (gray dots), n= 8 mice; *Scn8a*^+/mut^ (red dots), n=6 mice,; *Scn8a*^+/mut^ OPC cKO (blue dots), n=3 mice. 4-month-old time-point: *Scn8a*+/+ (black dots), n=5 mice; *Scn8a*^+/+^ OPC cKO (gray dots), n= 6 mice; *Scn8a*^+/mut^ (red dots), n=6 mice, *Scn8a*^+/mut^ OPC cKO (blue dots), n=4 mice. 6-month time-point: *Scn8a*+/+ (black dots), n=5 mice; *Scn8a*^+/+^ OPC cKO (gray dots), n=5 mice; *Scn8a*^+/mut^ (red dots), n=5 mice,; *Scn8a*^+/mut^ OPC cKO (blue dots), n=4 mice. Data represent mean ± SEM and data within each timepoint were analyzed with ANOVA followed by Tukey-Kramer test, adjusting for multiple comparisons. *p<0.05, **p<0.01, ***p<0.001, n.s. = p>0.05.

We next assessed whether deletion of the TrkB receptor from OPCs prevents the oligodendroglial response to seizures. Consistent with prior studies (Geraghty et al., 2019), OPC-specific deletion of TrkB does not alter homeostatic OPC numbers. Deletion of the TrkB receptor from OPCs in *Scn8a*^*+/mut*^;*TrkB*^*fl/fl*^; *PDGFRα::Cre* mice (*Scn8a*^*+/mut*^ OPC cKO) prevented the aberrantly increased callosal OPC number found in *Scn8a*^*+/mut*^ mice with intact activity-regulated myelin response, (**Figure 6B)**.

Having elucidated the timeline of epileptogenesis and confirmed that TrkB deletion from OPCs prevents OPC expansion in association with seizures, we next examined seizure frequency in *Scn8a*^*+/mut*^ mice lacking TrkB expression in OPCs (*Scn8a*^*+/mut*^ OPC cKO). We found that seizure burden was strikingly reduced in *Scn8a*^*+/mut*^ OPC cKO mice with impaired activity-dependent myelination. *Scn8a*^*+/mut*^ mice with intact activity-regulated myelination exhibit a marked increase in the number of seizures per hour over the period of epileptogenesis (**Figure 6D-E**). In contrast, *Scn8a*^*+/mut*^ mice with OPC-specific loss of TrkB expression that lack activity-regulated myelination (**Figure 6B** and Geraghty et al., 2019) exhibit substantially fewer seizures per hour, an effect that was sustained at least until 6 months of age (**Figure 6E**). The mean duration of individual seizures in *Scn8a*^+/mut^ mice was 2.3 ± 0.2 seconds, consistent with previously published findings in *Scn8a*^+/mut^ mice (Makinson et al., 2017); the mean duration of individual seizures was not significantly different in *Scn8a*^+/mut^ cKO mice (**Supplemental Figure 4)**. Taken together, these findings indicate that activity-dependent myelination contributes to kindling of absence seizures during epileptogenesis.

## Discussion

Childhood absence epilepsy, historically called “petit mal” seizures, is a common genetic generalized epilepsy syndrome (Matricardi et al., 2014; Brigo et al., 2018). Although childhood absence epilepsy has been considered relatively benign, this disease is associated with considerable cognitive co-morbidity and poor psychosocial functioning (Verrotti et al., 2015; Shinnar et al., 2017). Moreover, up to 35% of cases are refractory to medical therapy for unknown reasons (Wirrell et al., 1996) and in 20 to 50% of patients, seizures return following medication withdrawal (Matricardi et al., 2014). The natural history of the disease involves rapid, progressive increases in seizure frequency and severity, similar to what is seen in animal models (Brigo et al., 2018). Recent evidence indicates that this period of seizure “kindling” is a key determinant of disease severity in humans as well as rodent models, such that early blockade of absence seizures and/or their downstream effects mitigates morbidity (van Luijtelaar et al., 2013; Pitkanen et al., 2015; Morse et al., 2019). It should also be noted that multiple (often medically intractable) epilepsy syndromes involve typical and atypical absence seizures, at times in combination with other seizures types, such as other generalized epilepsies and the often devastating epileptic encephalopathy Lennox-Gastaut syndrome (Arzimanoglou et al., 2009; Pack, 2019). Thus, there is an urgent need to understand mechanisms underlying epileptogenesis, which will be necessary for disease-modifying and/or curative treatments for absence and other forms epilepsy.

Given recent appreciation that activity-regulated myelination can influence neural network function (Pajevic et al., 2014; Noori et al., 2020; Steadman et al., 2020), we reasoned that excessive and aberrant neuronal activity might abnormally increase myelination within seizure networks in disorders such as absence epilepsy. Maladaptive myelination may, in turn, contribute to disease pathogenesis, including seizure kindling. Further supporting the hypothesis that increased myelination, particularly within corpus callosum, might contribute to seizure susceptibility or severity, a rat strain with a propensity for provoked seizures exhibits increased callosal myelination (Sharma et al., 2017). Here, in two distinct rodent models with spontaneous absence seizures and well-defined periods of epileptogenesis, we found increased oligodendrogenesis and myelination of the seizure network only after seizure onset. Increased myelination did not occur when seizures were pharmacologically treated. We observed both an increase in mean myelin sheath thickness as well as an increase in the number of myelinated corpus callosum axons. Whether the change in myelinated axon number reflects de novo myelination of previously unmyelinated axons or discontinuously myelinated axon segments (Tomassy et al., 2014) remains to be determined. Furthermore, it is not yet clear whether the increased sheath thickness reflects newly generated internodes or activity-regulated remodeling by existing oligodendrocytes (Swire et al., 2019). Finally, the effects of seizure-associated myelination on inhibitory interneurons (Zonouzi et al., 2019) and other neuronal subtypes remains to be explored in future studies. While these mechanistic questions remain to be elucidated, a role for activity-regulated myelination in absence epileptogenesis is clear. Genetic blockade of activity-dependent myelination during epileptogenesis reduced the frequency of daily seizures over time. Together, our findings indicate that maladaptive, activity-regulated myelination contributes to progressive increases (kindling) of absence seizures.

How might activity-regulated myelination become maladaptive, contributing to further disease progression? In general, epileptogenesis in absence and other forms of epilepsy is thought to reflect increased neuronal excitation and synchrony of neuronal firing (McCormick and Contreras, 2001; Jefferys et al., 2012; Pitkanen et al., 2015; Fogerson and Huguenard, 2016). Activity-regulated myelination promotes oscillatory synchrony (Pajevic et al., 2014; Noori et al., 2020; Steadman et al., 2020); thus, aberrantly increased activity-regulated myelination may contribute to thalamocortical hypersynchrony underlying absence epilepsy. Consistent with our observations of increased callosal myelination, abnormally increased interhemispheric synchrony between the somatosensory cortices is well demonstrated in absence epilepsy (Mishra et al., 2013). Additionally, changes in myelination and consequent alterations to temporal dynamics within the seizure network could influence spike timing-dependent synaptic plasticity (Bi and Poo, 2001) and thus affect neuronal excitation. Increased myelination might also serve as a compensatory mechanism that provides metabolic support and enables rapid firing during seizures (Nave, 2010; Funfschilling et al., 2012). Finally, oligodendroglial cells can influence neuronal excitability. Satellite oligodendrocytes are electrically coupled with astrocytes via gap junctions in a “syncytium” which buffers potassium to constrain neuronal excitability (Battefeld et al., 2016). Similarly, conditional deletion of the inward rectifying potassium channel Kir4.1 from oligodendrocytes impairs potassium clearance, leading to hyperexcitability and decreased seizure threshold (Larson et al., 2018). A subset of CNS oligodendrocytes express glutamine synthetase and directly modulate glutamatergic excitatory neurotransmission (Xin et al., 2019). In absence epilepsy, the impact of the observed increase in oligodendrogenesis and myelination upon potassium buffering and potassium-related neuronal excitability remains to be determined. Future work will investigate network level mechanisms by which myelin plasticity contributes to kindling, which may include promoting the thalamocortical and interhemispheric hyper-synchrony characteristic of absence epilepsy (Huntsman et al., 1999; Makinson et al., 2017).

Our finding that seizures are markedly reduced but not entirely prevented by blockade of activity-dependent myelination suggests that multiple mechanisms are responsible for progressive increases in daily seizure burden observed in *Scn8a* mice. This is consistent with previous findings that *Scn8a* loss of function, presumably unaffected by genetic blockade of activity-regulated oligodendrogenesis, leads to thalamocortical hypersynchrony and seizures due to interneuron dysfunction in the reticular thalamic nucleus (Makinson et al., 2017). Altered function of voltage-gated calcium channels (Pietrobon, 2002), GABA receptors (Seo and Leitch, 2014), hyperpolarization-activated cyclic nucleotide-gated potassium channels (Ludwig et al., 2003), and glucose transport (Marin-Valencia et al., 2012) also cause or contribute to absence epileptogenesis. Extensive studies have elucidated altered neuronal physiology in the thalamocortical network leading to absence seizures (von Krosigk et al., 1993; Steriade and Contreras, 1998; Huntsman et al., 1999; Paz et al., 2011), while the involvement of glial cells in the pathogenesis of absence epilepsy is only beginning to be recognized. The early studies of glia in epilepsy pathogenesis have focused chiefly on astrocytes (Coulter and Steinhauser, 2015). Impaired GABA transport in astrocytes was found to promote tonic, rather than phasic, GABA-A signaling, potentiating absence seizures in multiple models of absence epilepsy (Cope et al., 2009); astrocyte-mediated glutamate metabolism also regulates the duration of thalamocortical epileptiform oscillations (Bryant et al., 2009); endozepine modulation of GABA-A currents in the reticular thalamic nucleus, a significant determinant of seizure severity, was also found to be regulated by astrocytes (Christian et al., 2013; Christian and Huguenard, 2013). Increased GFAP expression has been noted throughout the thalamocortical network in absence epilepsy, further supporting the idea of astrocyte dysregulation or reactivity (Cavdar et al., 2019). More broadly, in multiple forms of epilepsy, an array of astrocyte-mediated mechanisms – including impaired glutamate and excitatory amino acid metabolism, potassium buffering, gap junction and aquaporin expression – are thought to contribute to hyperexcitability and seizures (Eid et al., 2019). Further, neuro-inflammation involving microglia, astrocytes and brain vasculature significantly modulates disease severity in some forms of epilepsy, reviewed in (Vezzani et al., 2019). In addition to these findings demonstrating effects of glia on neuronal hyperexcitability in multiple forms of epilepsy, here we hypothesize a role for oligodendrocytes in modulating neural network synchrony.

Given the diverse array of mechanisms occurring in different forms of epilepsy, it is likely that the role of myelin plasticity may also vary. For example, different forms of epilepsy involve distinct brain networks and different neuronal populations exhibit variable potential for activity-regulated myelination (Gibson et al., 2014). Furthermore, different seizure types may involve neurons firing at different rates (Steriade et al., 1998; Truccolo et al., 2011; Schevon et al., 2012), and some neurons in the same network may fire at increased or decreased rates (Truccolo et al., 2011); myelin-forming cells may be differentially affected by different neuronal firing rates (Stevens et al., 1998; Nagy et al., 2017). Thus, ictal and interictal patterns of neuronal activity may drive divergent patterns of myelination in affected and non-affected networks. Forms of epilepsy in which neurodegenerative and inflammatory mechanisms are particularly prominent, such as mesial temporal lobe epilepsy (Vezzani et al., 2011), might be associated with diminished white matter plasticity due to effects of reactive microglia on oligodendroglial cells (Miron et al., 2013; Gibson et al., 2019). Underscoring the likely heterogeneity of myelin changes in different seizure types, brief acute generalized tonic-clonic seizures did not induce an oligodendroglial response in a previous study (Gibson et al., 2014). Thus, future work should investigate the likely heterogeneous patterns and functional roles of myelination in diverse forms of epilepsy.

The findings presented here highlight avenues for potential therapeutic interventions targeting aberrantly increased oligodendrogenesis and myelination. In the case of absence seizures, we found that genetic blockade of activity-dependent myelination by targeted removal of TrkB from OPCs (Geraghty et al., 2019) reduced seizure burden. Therapeutically targeting BDNF signaling, a pathway critical to many adaptive processes (Kowianski et al., 2018) may confer risks to cognition and neurodevelopment that outweigh the benefit to seizure severity. Alternatively, oligodendrogenesis can be targeted using pharmacological histone deacetylase (HDAC) inhibitors to epigenetically interrupt oligodendroglial differentiation (Wu et al., 2012; Gibson et al., 2014). Indeed, HDAC inhibition has been shown to improve the course of absence epilepsy in Wag/Rij rats (Citraro et al., 2020), although the link to myelination has not been previously appreciated. Such strategies targeting aberrant oligodendrogenesis may prove particularly helpful in refractory cases of childhood absence epilepsy, and/or in preventing oligodendrogenesis in epilepsy syndromes defined by intractable forms of absence seizures, such as Lennox-Gastaut syndrome (Camfield, 2011). More broadly, the finding that aberrant activity-regulated myelination can contribute to seizure kindling suggests that maladaptive myelination may be a therapeutically targetable pathogenic mechanism in neurological and neuropsychiatric diseases defined by recurrent patterns of abnormal neuronal activity.

## Author Contributions

J.K.K. performed experiments and analyzed quantitative microscopy and electrophysiological data. C.S., E.F., L.T.T., D.F., A.B., T.S. and H.X. performed experiments and assisted with data analysis. L.T.T., A.B. and K.V. assisted with animal husbandry and drug administration. L.N. performed electron microscopy. J.K.K., M.M. and J.H. conceived of the project. J.K.K., M.M. wrote the manuscript. H.X., A.B., and E.F. C.S., E.F., L.T.T., D.F., A.B., T.S. and H.X. edited the manuscript. M.M. and J.H. supervised all aspects of the work.

## Acknowledgements

The authors gratefully acknowledge support from the National Institute of Neurological Disorders and Stroke (R01NS092597 to M.M, K12NS098482-02 to J.K.K., R01NS034774 to J.H.), NIH Director’s Pioneer Award (DP1NS111132 to M.M.), Kleberg Foundation, Stanford Maternal and Child Health Research Institute (to M.M. and J.K.K.), Bio-X Institute (to M.M. and J.K.K.), Cancer Research UK (to M.M.), American Epilepsy Society and CURE Epilepsy Foundation (to J.K.K.). The authors wish to thank Michelle Fogerson, Jordan Sorokin, Austin Reese and Christopher Makinson for their guidance on performing and analyzing rodent EEG. The authors also thank Dr. Steve Chinn at Stanford Children’s Health for his assistance with ethosuximide experiments.

---

**Supplemental Figure 1:**
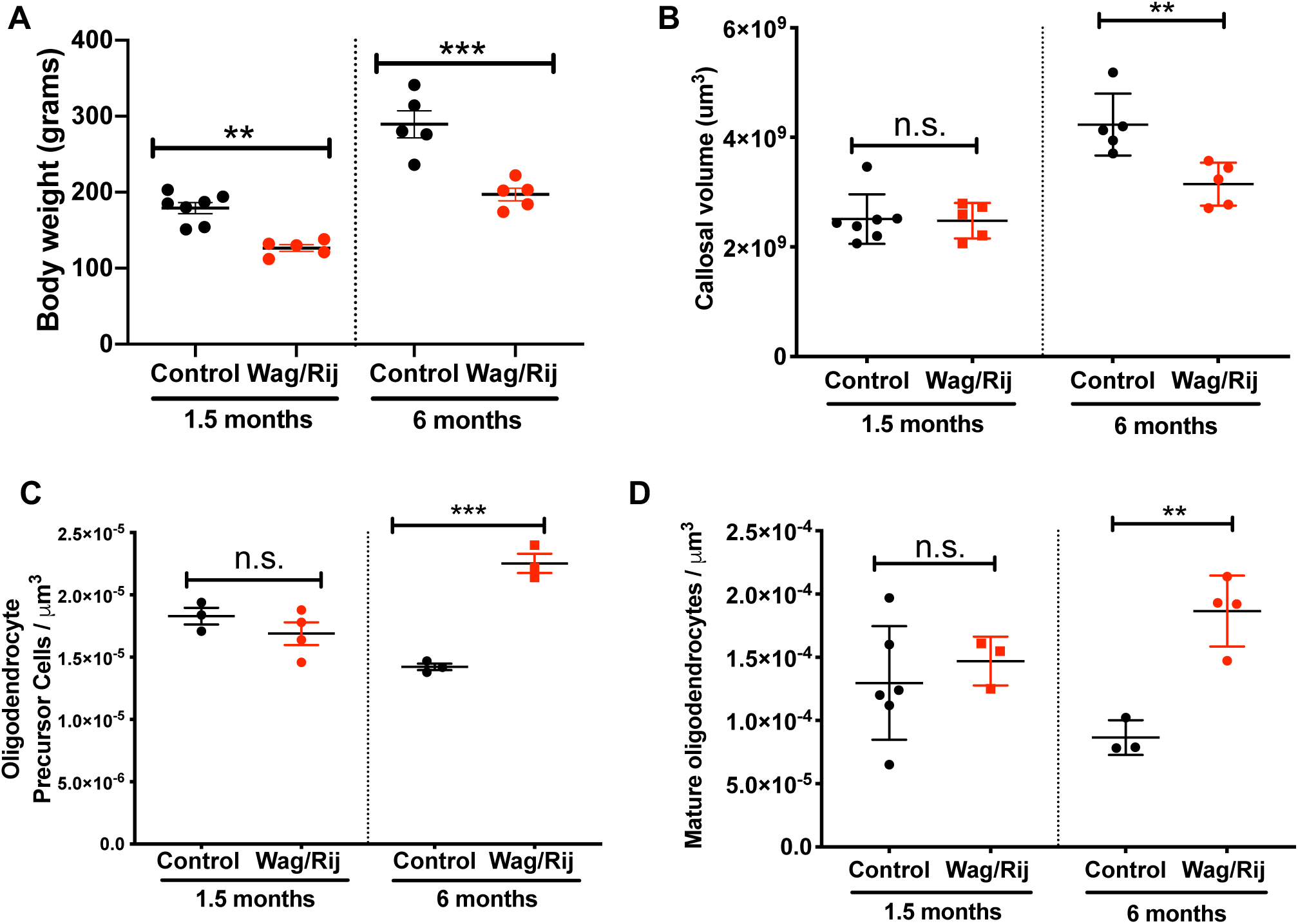
Oligodendroglial density increases with absence seizures in Wag/Rij rats. **Related to Figure 1** **(A)** Control (Wistar; black dots) and Wag/Rij (red dots) rats were weighed at 1.5 and 6 months of age. Each dot is the weight in grams for one animal; data represent mean ± SEM. Data were analyzed by ANOVA followed by Sidak’s post-hoc testing (1.5-month timepoint: control =7 rats; Wag/Rij =5. 6-month timepoint: control = 5 rats, Wag/Rij = 5). **(B)** The volume of the body of the corpus callosum was computed with Cavalieri’s method in control (black dots) and Wag/Rij rats (red dots). Each dot is the callosal volume for one rat; data represent mean ± SEM. Data were analyzed by ANOVA followed by Sidak’s post-hoc testing. (1.5-month timepoint: control =7 rats; Wag/Rij =5. 6-month timepoint: control = 5 rats, Wag/Rij = 5). **(C)** Unbiased stereological assessment of OPC density (OPC number normalized to callosal volume). OPCs were defined as cells expressing PDGFR*α* and Olig2. Each dot is the OPC density for one mouse; data represent mean ± SEM. Data were analyzed by ANOVA followed by Sidak’s post-hoc testing. (1.5-month timepoint: control =3 rats; Wag/Rij =4. 6-month timepoint: control = 3 rats, Wag/Rij = 3). **(D)** Unbiased stereological assessment of mature oligodendrocyte density. Mature oligodendrocytes expressed Olig2 and CC1 but not PDGFR*α*. Each dot is the oligodendrocyte density for one rat; data represent mean ± SEM. Data were analyzed by ANOVA followed by Sidak’s post-hoc testing. (1.5-month timepoint: control = 6 rats; Wag/Rij = 3. 6-month timepoint: control = 3 rats, Wag/Rij = 4).

**Supplemental Figure 2:**
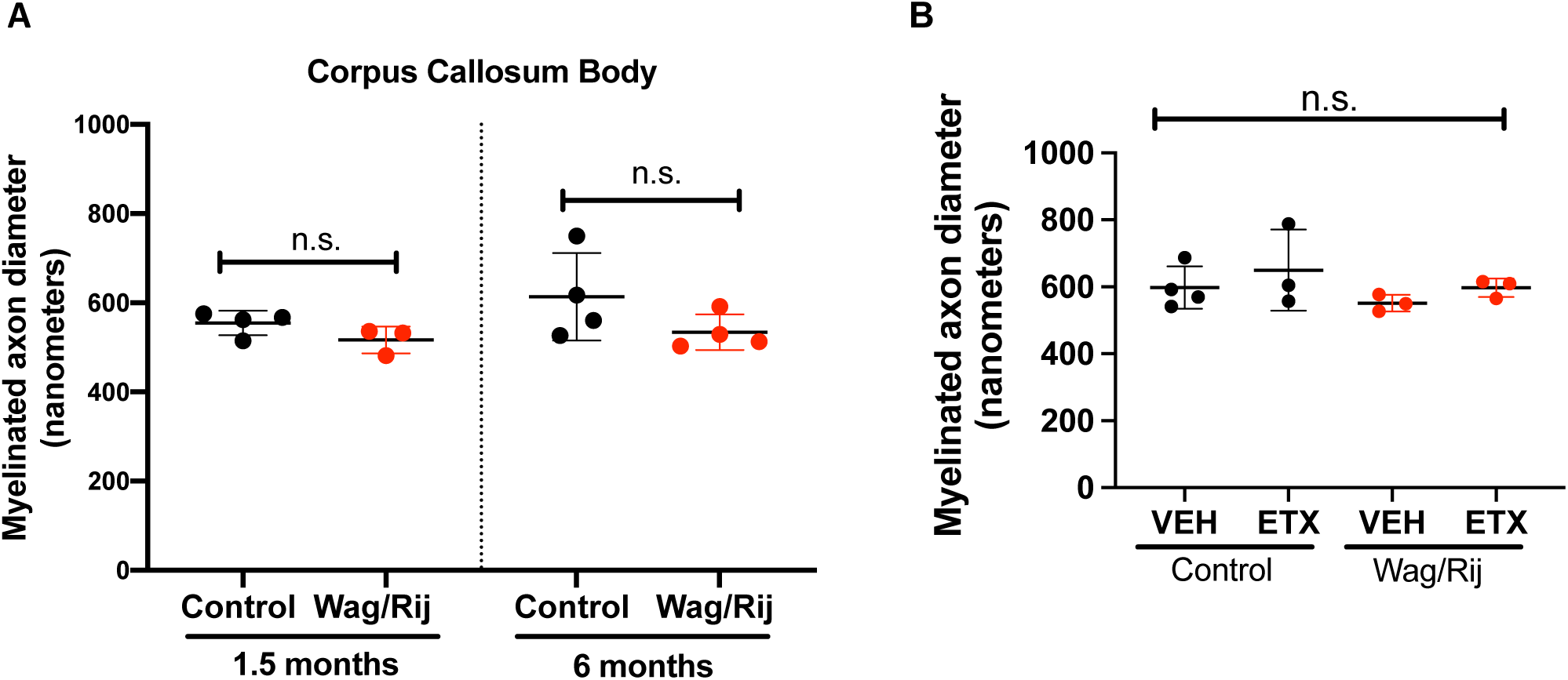
Myelinated axon diameters do not contribute to *g*-ratio differences in Wag/Rij and control rats. **Related to Figure 2**. **(A)** Mean myelinated axon diameter in the body of the corpus callosum in control (black dots) and Wag/Rij (red dots) rats at 1.5 and 6 months of age. 1.5 month timepoint, n = 4 control, 3 Wag/Rij; 6 month timepoint, n = 4 control, 4 Wag/Rij rats. Data were analyzed with ANOVA with post-hoc Sidak’s testing and represent mean ± SEM; each dot represents one animal. **(B)** Control and Wag/Rij rats exhibit similar axon diameter, with and without ethosuximide (ETX) treatment. Mean myelinated axon diameter at 7 months of age in control (black dots) and Wag/Rij (red dots) rats treated with vehicle (VEH) or ETX. (Control-VEH, n = 4; Control-ETX, n = 3; Wag/Rij-VEH, n = 3; Wag/Rij-ETX, n = 3). Data represent mean ± SEM and were analyzed by ANOVA with Tukey-Kramer post-hoc testing.

**Supplemental Figure 3:**
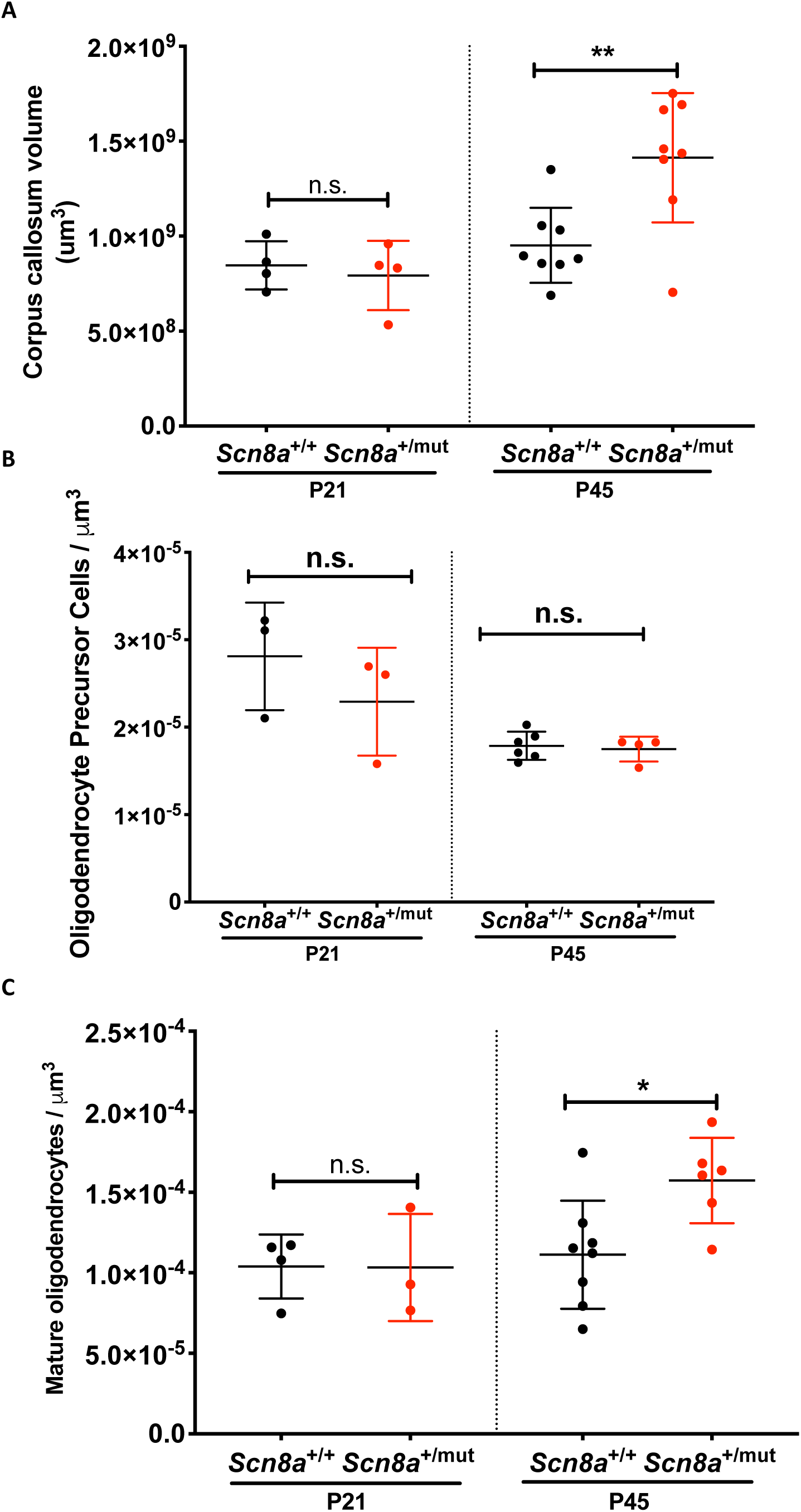
Increased oligodendroglial density in the absence seizure network of *Scn8a*^+/mut^ mice. **Related to Figure 4**. **(A)** The volume of the body of the corpus callosum was computed with Cavalieri’s method in *Scn8a*^+/+^ mice (black dots) and *Scn8a*^+/mut^ mice (red dots). Each dot is the callosal volume for one mouse; data represent mean ± SEM. (1.5-month timepoint: *Scn8a*^+/+^ =4 mice; *Scn8a*^+/mut^ = 4 mice. Six-month timepoint: *Scn8a*^+/+^ = 8 mice, *Scn8a*^+/mut^ = 8 mice). **(B)** Unbiased stereological quantification of OPC density (cells co-expressing PDGFR*α* and Olig2) in the body of the corpus callosum P21 (prior to seizure onset) and P45 (after seizures are well-established in *Scn8a*^+/mut^ mice). (P21, *Scn8a*^+/+^ n= 3; *Scn8a*^+/mut^ n = 3. P45, *Scn8a*^+/+^ n= 6; *Scn8a*^+/mut^ n = 4). Black dots represent wildtype littermates (*Scn8a*^+/+^) while red dots represent *Scn8a*^+/mut^ mice. Data represent mean ± SEM; each dot represents the OPC density for one animal. **(C)** Unbiased stereological quantification of mature oligodendrocyte density (cells co-expressing CC1 and Olig2) in the body of the corpus callosum of *Scn8a*^+/+^ (black dots) and *Scn8a*^+/mut^ mice (red dots) at P21 and P45. (P21, *Scn8a*^+/+^ n= 4; *Scn8a*^+/mut^ n = 3. P45, *Scn8a*^+/+^ n= 8; *Scn8a*^+/mut^ n = 6). Data represent mean ± SEM; each dot represents the oligodendrocyte density of one animal. For all panels in this figure, data were analyzed with ANOVA with post-hoc Sidak’s testing with correction for multiple comparisons. *p<0.05, **p<0.01, ***p<0.001, n.s. = p>0.05.

**Supplemental Figure 4:**
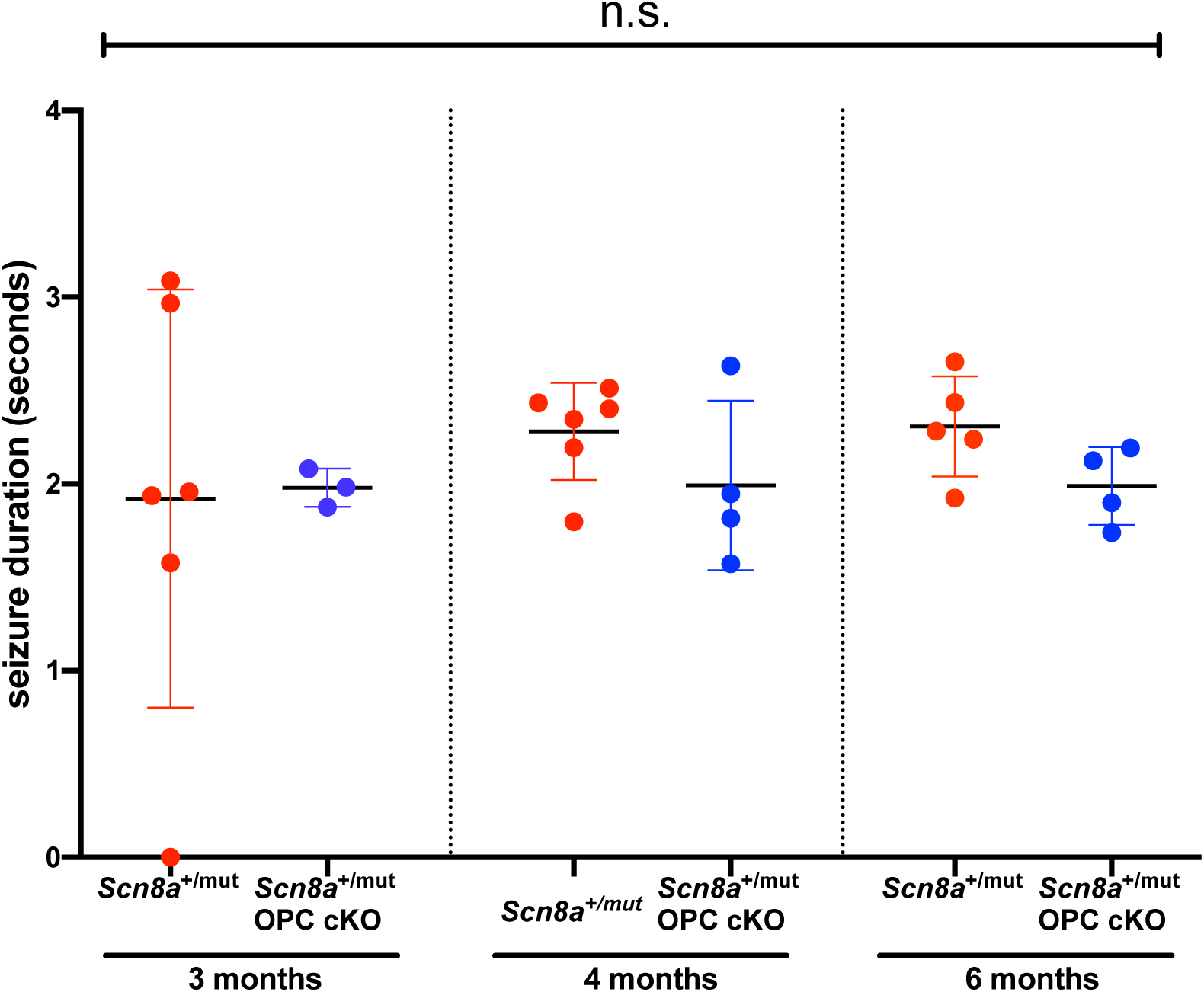
Activity-dependent myelination does not impact seizure duration. Related to Figure 6. Mean seizure duration was quantified in *Scn8a*^+/mut^ (red dots) and *Scn8a*^+/mut^ OPC cKO mice (blue dots). 3-month-old time-point: *Scn8a*^+/mut^ n=6 mice, *Scn8a*^+/mut^ OPC cKO, n=3 mice. 4-month-old time-point: *Scn8a*^+/mut^, n=6 mice, *Scn8a*^+/mut^ OPC cKO, n=4 mice. 6-month time-point: *Scn8a*^+/mut^, n=5 mice, *Scn8a*^+/mut^ OPC cKO, n=4 mice. Data represent mean ± SEM and data were analyzed with ANOVA followed by Sidak’s test.

## STAR Methods

### LEAD CONTACT

Further information and requests for resources and reagents should be directed to and will be fulfilled by the Lead Contact, Michelle Monje.

## MATERIALS AVAILABILITY

- This study did not generate new unique reagents.

## DATA AND CODE AVAILABILITY

- The MATLAB code used for EEG analysis in this study is available from the corresponding authors on request.
- Data will be available on Mendeley in the final version of the manuscript and are also available upon request now.

## EXPERIMENTAL MODEL AND SUBJECT DETAILS

### Rodent colony maintenance

All experiments were conducted in accordance with protocols approved by the Stanford University Institutional Animal Care and Use Committee (IACUC). Mice or rats were group or single housed (up to 5 mice or 2 rats per cage) according to standard guidelines with *ad libitum* access to food and water in a 12 h light/dark cycle. No animals were manipulated other than as reported for that experimental group, i.e., there was no history of drug exposures, surgeries or behavioral testing for the animals used other than that reported for the given experimental group. Mice and rats were healthy and tolerated all experimental manipulations well.

The ages of all mice and rats used in specific studies are indicated in the figures and throughout the text. Briefly, in studies using Wag/Rij and control rats, 1.5-month-old animals were used to assess endpoints prior to seizure onset and 6-7-month-old animals were used to assess endpoints after seizures are well established. In studies using *Scn8a*^+/mut^ mice and wild-type littermates, post-natal day (P)21 mice were used to assess endpoints prior to epileptogenesis, while p45 mice were used to assess endpoints after seizures are well established. In studies in which *Scn8a*^+/mut^ and other mouse lines were bred (e.g. Figure 6), because seizure onset is delayed, later time points (3 to 6 months) were used to study seizure progression, as described in detail in the text and methods below.

Both males and females (rats and mice) were used in equal numbers whenever possible. There was no impact of male or female sex upon any of the endpoints. For the majority of experiments, individual animals utilized came from ≥ 2 distinct litters.

### Wag/Rij rats

Wistar (control) and Wag/Rij rats were purchased from Charles Rivers Laboratories (Wistar: cat #003; Wag/Rij, Charles Rivers Italy: strain code #638). A colony of Wag/Rij rats has subsequently been maintained in Dr. Huguenard’s lab at Stanford (Sorokin et al., 2017).

### *Scn8a*^+/mut^ mice and *Scn8a*^+/mut^ OPC cKO mice

*Scn8a*^+/mut^ mice with the *med* loss of function mutation in *Scn8a* (C3Fe.Cg-Scn8amed/J, jax.org/strain/003798; Makinson et al, 2017) were bred to wild-type mice (*Scn8a*^+/+^) on a congenic background strain (C3HeB/FeJ, jax.org/strain/00658). In separate studies, successive breeding was performed to obtain mice with absence seizures (*Scn8a*^+/mut^) and inducible deletion of the TrkB receptor from OPCs, enabling blockade of activity-dependent myelination (*Scn8a*^+/mut^; *TrkB*^fl/fl^; *PDGFRα*::*Cre*-ER). Specifically, *Scn8a*^+/mut^ mice on the C3HeB/FeJ background were crossed with mice expressing a “floxed” *TrkB* gene (*TrkB*^fl/fl^) on a C57/BL6 background to obtain *Scn8a*^+/mut^ and *Scn8a*^+/+^ mice with floxed *TrkB* (*Scn8a*^+/mut^; *TrkB*^fl/fl^ or *Scn8a*^+/+^; *TrkB*^fl/fl^). These mice were crossed with mice with floxed *TrkB* and tamoxifen inducible *Cre* under the PDGFRa promotor (*TrkB*^fl/fl^; *PDGFRα*::*Cre*-ER), also on a C57/BL6 background. Both the *PDGFRα*::*Cre*-ER (originally acquired from The Jackson Laboratory, jax.org/strain/018280) and *TrkB*^fl/fl^ (originally acquired form MMRRC at UC Davis, strain # 033048-UCD) mouse lines are maintained in our laboratory, and are described in previously published work (Geraghty et al., 2019).

Mice were treated with Tamoxifen (Sigma-Aldrich, T5648) 100 mg/kg IP for 3 days (p21-p23) which induces robust *Cre* expression and deletion of the TrkB receptors from OPCs in *TrkB*^fl/fl^; *PDGFRα*::*Cre*-ER mice but not mice lacking *Cre* (*TrkB*^fl/fl^ only)(Geraghty et al., 2019).

### Genotyping

At P10, mice were ear and/or tail clipped and DNA was extracted using the REDExtract-N-Amp™ Tissue PCR Kit (Sigma-Aldrich, Cat# XNAT-100RXN). A master mix of REDEX reagent, primers as indicated below, water, and extracted DNA from each animal were created for a total volume of 20 µL for each animal. For detection of the floxed TrkB allele, the following primers (acquired from IDT) were used: TrkB fl/fl F: ATGTCGCCCTGGCTGAAGTG ; TrkB fl/fl R: ACTGACATCCGTAAGCCAGT. The PCR protocol was 94°C for 3 minutes ⨯ 1; then 94°C for 1 minute; 65°C for 1.5 minutes; 72°C for 1.5 minutes all x 40 cycles; then 72°C for 7 mins ⨯ 1. The PCR products were run on a 1.8 % agarose gel and a band size of 450 bp indicated the presence of the floxed allele. For detection of *Cre*, the following primers (IDT) were used: Cre Up: GAT CTC CGG TAT TGA AAC TCC AGC; Cre Down: GCT AAA CAT GCT TCA TCG TCG G. PCR protocol was 94°C for 5 minutes ⨯ 1; 94°C ⨯ 15 seconds, 65°C ⨯ 30 seconds, dropping 1°C after each cycle until the last cycle occurred at 55°C, all ⨯ 10 cycles; then 72°C for 40 seconds ⨯ 1 cycle. The presence of a 650 bp PCR product on a 1.8 % agarose gel indicated an animal positive for the *PDGFRα::Cre* gene. Automated genotyping for the *Scn8a*^med^ allele was done using tail samples via Transnetyx, Inc (www.transnetyx.com).

## METHOD DETAILS

### Ethosuximide treatment paradigm

Wistar (control) and Wag/Rij rats were treated with ethosuximide (ETX) or vehicle on a previously published dosing schedule (Blumenfeld et al., 2008) shown to prevent and/or significantly reduce seizures. Male and female littermates were randomly assigned to either vehicle or ETX treatment. Pharmaceutical grade ETX solution (Akorn Pharmaceuticals, 250 mg/mL, NDC: 61748-024-16) was added to drinking water in bottles that were shielded from light, at a concentration of 2.5 mg/mL leading to approximately dosing of 300mg/kg/day dosing (Blumenfeld et al., 2008). The solution was changed at least every 7 days and the health, weight and drinking of animals was monitored regularly throughout the study. Prior to perfusion, 1-2 mL of blood was taken. The samples were centrifuged at 2000rpm, 4 degrees C, for 5 minutes and plasma was collected for measurement of ETX concentration using liquid chromatography/mass spectroscopy, performed at the Stanford BioADD center (http://med.stanford.edu/bioadd.html).

### *In vivo* Electrocorticography (EEG)

Rats were stereotactically implanted with wires attached to screws over the bilateral somatosensory cortices (AP -3mm, ML 3.5-4mm), at 6 months of age. Mice were implanted for EEG following tamoxifen administration. *Scn8a*^+/mut^; *TrkB*^fl/fl^; *PDGFRa*::*Cre*-ER mice and their littermates were implanted for EEG with wires (A-M Systems, Cat#791400) and screws (J.I. Morris, Cat#FF00CE125) over the bilateral somatosensory cortices (coordinates AP -1.3 mm, ML 1.5-2 mm).

Mice and rats were also implanted with a reference wire over cerebellum, and implants were secured with dental cement (Metabond, #S399, S371, S398; also Jet Set4 Liquid, Lang Dental 3802×6). Implanted wires were integrated into custom-made Mill-Max headpieces (Digi-Key Electronics, ED90267-ND) that could be connected to a head stage, consisting of a digitizer and amplifier board (Intan Technologies, C3334). Awake and freely behaving animals were tethered to an acquisition board (Open Ephys) with light-weight SPI interface cables (Intan, C3206). Continuous real-time EEG was recorded with Open Ephys software (https://open-ephys.org). Data were sampled at 2 kHz and bandpass filtered between 1-300Hz. All animals underwent 3-4 hours of continuous EEG recording between the hours of 10am-5pm.

### Immunohistochemistry

Rats were given a lethal dose of Fatal Plus (sodium pentobarbital, Vortech Pharmaceuticals, NDC: 0298-9373-68) and transcardially perfused with 40-80 mL of ice-cold 0.1 M PBS followed by 40-80 mL 4% paraformaldehyde in PBS before brain extraction and tissue processing. Mice were given a lethal dose of Fatal Plus or 2.5% Avertin (2,2,2-tribromoethanol, Sigma, T48402) and transcardially perfused with 10-20 mL of ice-cold PBS followed by 10-20 mL of 4% paraformaldehyde in PBS. In studies involving ETX-treated rats and *Scn8a*^+/mut^; *TrkB*^fl/fl^; *PDGFRa*::*Cre*-ER mice, brains were bisected in the sagittal plane at the interhemispheric fissure. One half was post-fixed in 4% paraformaldehyde overnight at 4C prior to transfer to 30% sucrose for cryoprotection for several days; the other half was processed for electron microscopy (see below). Cryoprotected brains or hemi-brains were embedded in Tissue-Tek OCT compound (Sakura, 4583) and sectioned in the coronal plane at 50 μm using a sliding microtome (Microm HM450; Thermo Scientific). For immunohistochemistry, selected coronal sections were rinsed three times in TBS and incubated in blocking solution (3% normal donkey serum, Jackson Immunoresearch AB2337258; 0.3% Triton X-100 in TBS) at room temperature for 30 min. Rabbit anti-Olig2 (1:400, Millipore AB9610), Goat anti-PDGFRα (1:200, R&D Systems AF1062), mouse anti-APC (CC1,1:50, Calbiochem, OP80), or Rat anti-Ki67 (1:200, Life Technologies, 14-5698-82) were diluted in staining solution (1% normal donkey serum, 0.3% Triton X-100 in TBS) and incubated with sections at room temperature for 2.5 days; when staining for CC1, sections were incubated at 4C for 10 days. Sections were then rinsed three times in TBS and incubated in secondary antibody solution containing Alexa 488 donkey anti-rabbit IgG (1:500, Jackson Immuno Research, 711-545-152), Alexa 647 donkey anti-goat IgG (1:500, Jackson Immuno Research, A21447), Alexa 594 donkey anti-mouse IgG (1:500, Jackson Immuno Research, 715-585-150), or Alexa 594 donkey anti-rat IgG (1:500, Jackson Immuno Research, 712-585-153) in staining solution at 4°C overnight. Sections were rinsed three times in TBS and mounted with ProLong Gold Mounting medium (Life Technologies, P36930).

### Electron microscopy

Rats were given a lethal dose of Fatal Plus (sodium pentobarbital, Vortech Pharmaceuticals 0298-9373-68) and transcardially perfused with 40-80 mL of ice-cold 0.1 M PBS followed by 40-80 mL Karnofsky fixative, consisting of 4% PFA (Electron Microscopy Services (EMS), 15700) and 2% glutaraldehyde (EMS, 16000) in 0.1M sodium cacodylate buffer (EMS, 12300) (Geraghty et al., 2019; Gibson and Monje, 2019; Gibson et al., 2014) in PBS before brain extraction and tissue processing. Mice were given a lethal dose of Fatal Plus or Avertin (Sigma, T4,8402) and transcardially perfused with 10-20 mL of ice-cold PBS followed by 10-20 mL of Karnofsky fixative. In experiments where hemi-brains were used, one half was post-fixed in Karnofsky’s fixative immediately following perfusion with PBS and 4% PFA. Samples were post-fixed in Karnofsky fixative for at least 2 weeks. A ∼1mm^3^ block of tissue was dissected from the midline sagittal corpus callosum at the rostrocaudal location overlying the dentate gyrus and the hippocampal fornix, enabling cross-sectional views of callosal projection axons. The dissected block was processed for transmission electron microscopy as described previously (Gibson et al., 2014). Briefly, tissue was post-fixed in 1% osmium tetroxide (EMS 19100) for 1 hour at room temperature, washed three times in ultra-filtered water, then en bloc stained for 2 hours at room temperature before dehydration in gradient ethanols, then rinsed in 100% ethanol twice followed by acetonitrile (Fisher Scientific, A21-1). Samples were then embedded in 1:1 Embed-812 (EMS 14120):acetonitrile, followed by EMbed-812 for 2 hours, then placed into TAAB capsules filled with fresh resin before incubation in a 65°C oven overnight. 75-90nm sections from this block were mounted on Formvar-carbon coated slot grids and contrast stained for 30 seconds in 3.5% uranyl acetate in 50% acetone (EMS, 10015), followed by 0.2% lead citrate (EMS, 0378) for 30 seconds. Samples were imaged with a JEOL JEM-1400 transmission electron microscope at 120kV and images were collected with a Gaton Orius digital camera.

## QUANTIFICATION AND STATISTICAL ANALYSIS

### Unbiased stereology

All quantifications were performed by experimenters blinded to subject identity and experimental condition. OPCs and oligodendrocytes were visualized with an MBF Zeiss Axiocam light microscope. Cell numbers were determined through unbiased stereology using Stereo Investigator software (MBF Bioscience). Regions of interest containing the body of the corpus callosum, beginning rostrally at the level of the hippocampal fornix at the third ventricle, and continuing caudally through the anterior hippocampus (approximate Bregma AP –0.8 mm to –3.8 mm for rats, -0.8mm to –2 mm from mice) were traced within sections at 2.5x magnification. Images were acquired at 63x from every 6^th^ section throughout this region; 40x magnification was used for analyses of Ki67-expressing OPCs. For mice, this included 3-6 sections per animal, while for rats, this included 4-7 sections per animal; there were no differences in the number of sections used between experimental groups. Stereological parameters were determined through pilot studies, ensuring that at least 100-300 cells would be counted per animal and the Gunderson m=1 coefficient of error was less than 0.1 (Slomianka and West, 2005; West, 2013) (typical CEs were 0.03-0.07). In all studies, disector height of 30 mm and guard zones of 3 mm were applied unless otherwise indicated. We measured the volume of the body of the corpus callosum (region interrogated for cell counts) with the Cavalieri method (West, 2012). The following stereological parameters were used:

- Mouse OPCs: sampling grid size was set to 225 x 225 μm; counting frame was set to 100 ⨯ 100 μm.
- Mouse oligodendrocytes: sampling grid: 500 ⨯ 250 μm; counting frame 100 ⨯ 100 μm, disector height 20 μm.
- Rat OPCs: sampling grid size, 300 ⨯ 150 μm; counting frame 75 x 75 μm.
- Rat OPCs in hemibrains (ETX studies): grid size 225 ⨯ 225 μm; counting frame 100 x 100 μm.
- Rat Ki67-expressing OPCs: 300 x 175 μm; counting frame 130 ⨯ 130 μm.
- Rat oligodendrocytes: 750 x 350 μm; counting frame 75 ⨯ 75 μm; disector height 20 μm.

### Quantitative analysis of myelin structure

*G*-ratio, defined as the length of the axonal diameter in its short axis, divided by the diameter of the entire fiber in the same axis (axonal diameter / axonal diameter + myelin sheath), was quantified using ImageJ software (https://imagej.nih.gov/ij/) (Gibson et al., 2014; Steadman et al., 2020; Waxman, 1980). Axons were quantified from 4000X transmission electron micrographs. Myelinated axon number was determined by quantifying the number of myelinated axons per 4000x electron micrograph and multiplying by the measured volume (in μm ^3^) of the corresponding region (body) of the corpus callosum. ∼150-300 axons from 8-20 micrographs were quantified per animal (exact ranges are specified in figure legends).

### EEG analysis

EEG data acquired with Open Ephys software was displayed and seizures from 3-4 hours of EEG recording were visually identified, marked and tabulated by a blinded reviewer using custom MATLAB software. Seizures were associated with behavioral arrest, and the corresponding EEG demonstrated predominantly 4-8 Hz frequency spike wave morphology, amplitude ∼1.5-2 times that of the background, and duration > 1 second (Makinson et al., 2017; Sorokin et al., 2017).

### Statistical analysis

Full details of statistical analyses can be found in the figure legends. For all studies, “n” refers to the number of mice or rats included in each experimental group, and unless indicated otherwise (e.g. g-ratio scatterplots), each data point in a graph represents the mean from one mouse. For all studies, n= 3 or more mice / rats per group, with the exact n specified in figure legends. Sample sizes were based on the variance of data in pilot experiments, and were generally estimated by power calculations which determined the number of animals “n” needed for 80% power to detect a 20-30% difference between genotypes. GraphPad Prism software (version 8, GraphPad Software, LLC) was used to perform statistical analyses. Statistical significance was defined as p < 0.05 throughout. The Wilks-Shapiro test was used to determine whether data were normally distributed; parametric tests were used where indicated. Nonparametric tests were used for nonparametric datasets. For experiments comparing control or Wag/Rij rats, or *Scn8a*^+/+^ and *Scn8a*^+/mut^ mice at different time points (e.g. Figures 1, 2, 4, 5 and Supplemental Figures 1, 2A, and 3) ANOVA followed by Sidak’s post hoc-test, correcting for multiple comparisons, was used to specifically compare groups within specific age groups. When two experimental groups within only one time-point were compared (Figure 1F), a t-test was used. For experiments in which rats were treated with ethosuximide or vehicle (Figure 3), and those studies involving conditional knockout of TrkB from OPCs in *Scn8a*^+/+^ and *Scn8a*^+/mut^ mice (Figure 6, Supplemental Figure 4), ANOVA with post-hoc Tukey-Kramer testing was employed in order to perform multiple relevant comparisons with appropriate corrections. For comparisons of seizure frequency in rats (Figure 3B), the nonparametric Kruskal-Wallis and Dunn’s post-hoc tests were used.

